# Evolutionary divergence between homologous X-Y chromosome genes shapes sex-biased biology

**DOI:** 10.1101/2024.03.27.586985

**Authors:** Alex R. DeCasien, Kathryn Tsai, Siyuan Liu, Adam Thomas, Armin Raznahan

## Abstract

Sex chromosomes are a fundamental aspect of sex-biased biology, but the extent to which homologous X–Y gene pairs (“the gametologs”) contribute to sex-biased phenotypes remains hotly-debated. Although these genes exhibit large sex differences in expression throughout the body (XX females express both X members; XY males express one X and one Y member), there is conflicting evidence regarding the degree of functional divergence between the X and Y gametologs. Here, we use co-expression fingerprint (CF) analysis to characterize functional divergence between the X and Y members of 17 gametolog gene pairs across >40 human tissues. Gametologs exhibit functional divergence between the sexes that is driven by divergence between the X vs. Y gametologs (assayed in males) and is greatest among evolutionary distant gametolog pairs. These patterns reflect that X vs. Y gametologs show coordinated patterns of asymmetric coupling with large sets of autosomal genes, which are enriched for functional pathways and gene sets implicated in sex-biased biology and disease. These findings suggest that the X and Y gametologs have diverged in function, and prioritize specific gametolog pairs for future targeted experimental studies.

## Introduction

Humans exhibit sex/gender* differences in diverse traits, including the prevalence and presentation of many medical conditions (*see Note 1) [1–3]. Two foundational biological sources for phenotypic sex differences in placental mammals (including humans) are: i) differences in circulating gonadal steroids and their downstream effects on diverse organ systems [4]; and ii) gonad-independent effects of differences in the dosage of X and Y chromosome genes [5]. Although mechanistic research on sex-biased biology has traditionally emphasized gonadal factors, there is growing recognition of the potential for sex chromosome genes to shape sex-biased traits [5]. Several lines of evidence suggest that a distinct subset of sex chromosome genes – the “gametologs” (Figures 1A, 1B) – may be key drivers of sex chromosome dosage effects on organismal structure and function.

**Figure 1.**
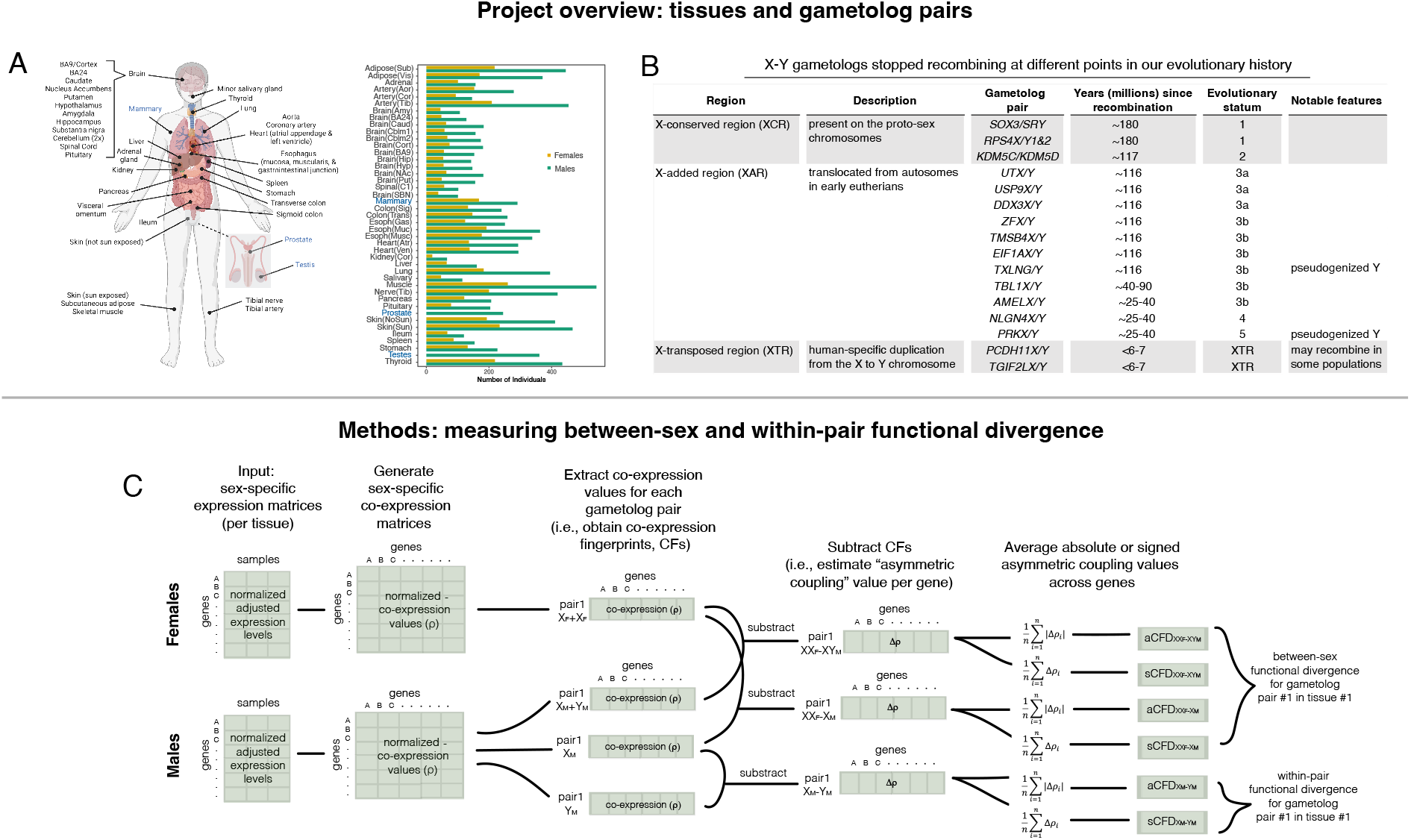
Overview of study design. a. Left: Tissues and samples included (cartoon is after GTEx [41] Figure 1). Created using BioRender.com. Right: Bar chart depicting the number of male (green) and female (yellow) samples available for each tissue. We verified robustness of asymmetric coupling after subsampling males to the minimum sample size across tissues (N = 66 in kidney) (Figure S6). Data are shown for N=43 tissues examined in this study (N=3 sex-specific or sex-differentiated tissues are in blue and are only included in male-specific analyses). BA9 = Brain(BA9); Cortex = Brain(Cort); BA24 = Brain(BA24); Caudate = Brain(Caud); Nucleus Accumbens = Brain(NAc); Putamen = Brain(Put); Hypothalamus = Brain(Hyp); Amygdala = Brain(Amy); Hippocampus = Brain(Hip); Substantia nigra = Brain(SBN); Cerebellum = Brain(Cblm1), Brain(Cblm2); Spinal Cord = Spinal(C1); Minor salivary gland = Salivary; Aorta = Artery(Aor); Coronary artery = Artery(Cor); Atrial appendage = Heart(Atr); Left ventricle = Heart(Ven); Esophagus mucosa = Esoph(Muc); Esophagus muscularis = Esoph(Musc); Esophagus gastrointestinal junction = Esoph(Gas); Kidney = Kidney(Cor); Transverse colon = Colon(Trans); Sigmoid colon = Colon(Sig); Adrenal gland = Adrenal; Visceral omentum = Adipose(Vis); Subcutaneous adipose = Adipose(Vis); Skin (not sun exposed) = Skin(NoSun); Skim (sun exposed) = Skin(Sun); Tibial nerve = Nerve(Tib); Tibial artery = Artery(Tib). b. Table listing gametolog gene pairs considered in this study. Evolutionary strata are from [91] and years since recombination are from [6]. Possible ongoing XTR recombination is noted by [43,44]. c. Analytic pipeline using co-expression fingerprints (CFs) to derive estimates of gene-level asymmetric coupling and of co-expression fingerprint divergence (CFD), the latter of which is a computational proxy for functional divergence between the X and Y members of gametolog pairs.

Gametologs are homologous gene pairs that have retained functional copies on both the X and Y chromosomes, despite residing within regions of these chromosomes that stopped recombining with each other as long as 180 million years ago [6]. Retention of functional Y copies distinguishes gametologs from all other non-recombining X–Y gene pairs, which lost Y member function through pseudogenization and gene loss, yielding the striking heteromorphism observed between modern X and Y chromosomes [6–9]. Y gametolog retention is believed to reflect extreme dosage sensitivity, since X–Y gametologs regulate transcription, translation, and protein stability [6,10–14]. However, given the high expression sensitivity of gametologs [15,16], these regulatory functions suggest that any functional divergences between X and Y gametologs could lead to widespread sex differences in genome function. This potential is elevated by the fact that, in most tissues, there is a categorical difference in expression potential of gametolog genes between XX and XY individuals: i) XX females can express two X members of each gametolog pair because X gametologs tend to escape X chromosome inactivation (XCI) [13,14,17,18]; and ii) XY males can express both X and Y members, as Y gametologs are some of the only Y chromosome genes expressed outside the testes [19,20]. Thus, understanding whether the X and Y gametologs have remained functionally equivalent over evolutionary time is crucial for understanding the biology of sex chromosomes and their potential to shape phenotypic sex differences.

Available studies have generated conflicting arguments regarding the degree of functional divergence between the X and Y members of gametolog gene pairs. Observations that support functional equivalence include: i) high levels of within-pair (i.e., X vs. Y member) sequence homology [19] and co-expression [20]; ii) the presence of sex-specific regulatory processes that maintain biallelic expression of gametologs in both sexes [14]; and iii) isolated experimental examples of functional homology (e.g., knockdown of *ZFX* or *ZFY* in human fibroblasts induces highly correlated effects on autosomal gene expression) [21]. However, there are also several theoretical and empirical observations that argue for functional divergence between the X and Y gametologs: i) their lack of recombination has exposed the Y members to male-specific selection pressures over millions of years [22]; ii) X vs. Y gametologs reside in different chromatin contexts, creating opportunities for regulatory divergence [23,24]; iii) some portions of coding sequences that have diverged between X and Y members change protein domains that are critical for their function [25,26]; iv) there are exceptions to the general pattern of high co-expression between the X and Y members [19,20,27]; v) the total (i.e., combined X and Y) expression level of many gametolog pairs is male-biased across human tissues, and this difference can translate to the protein level [10,17,20,28]; vi) experimental examples demonstrate functional differences between the protein products of specific X–Y gametolog pairs [21,25,26,29–33]; vii) many variants within X gametologs exhibit sex-biased allele frequencies [34–36], suggestive of sex-biased fitness effects; and viii) for some disease-linked gametolog pairs, there is direct evidence that loss of function mutations to the X vs. Y members do not cause equivalent effects [37,38].

We reasoned that resolving these tensions would require a systematic assay that can comprehensively and quantitatively index the degree of functional equivalence between X and Y gametologs across all pairs and tissues. Moreover, this assay should provide a path to biological annotation and analysis of disease relevance for any potential functional differences observed. Here, we utilize co-expression fingerprints (CFs) as scalable bioinformatic signatures of gene function. Because genes with highly correlated expression levels also share regulatory and functional relationships [39,40], testing for divergent CFs between X vs. Y gametologs offers an attractive computational assay for functional divergence. Crucially, analysis of CF divergence (CFD) can be systematically applied across gametolog pairs and tissues and also supports annotation of any gene sets that are differentially coupled to (i.e., co-expressed with) the X vs. Y gametologs. To this end, we: i) used measures of gene expression for >15k tissue samples (>800 individuals and >40 tissues from the GTEx resource [41]) (Table S1) to compute tissue-specific CFs for each X and Y gametolog gene; and ii) compared these CFs to derive tissue-specific estimates of functional divergence for each of 17 X–Y gametolog pairs (Figure 1, Methods) [7,9,14,42–44]. For each gametolog pair in each tissue, these CF comparisons can be applied in a sex-dependent manner (using CFs for the combined expression of both X members in females vs. the combined expression of the X and Y members in males) (Figure 1C) or in a sex-chromosome-dependent manner (using CFs for the X vs. Y member in males) (Figure 1C). The former measures sex differences in CFs (i.e., “between-sex CFD”) and the latter measures X–Y differences in CFs (i.e., “within-pair CFD”) (Figure 1C). Moreover, these divergence metrics can be estimated as absolute values (aCFD) to provide directionless measures of divergence, or as signed values (sCFD) that capture whether the rest of the transcriptome tends to be co-expressed more strongly with e.g., the X or Y member (i.e., positive vs. negative within-pair sCFD values, respectively). In the text below, we use the general term “functional divergence” and indicate the specific CFD metric being referred to in parentheses.

Our findings reveal that gametologs show high between-sex functional divergence (relative to other genes), and that this sex difference is driven by functional divergence between the X vs. Y gametologs (which is estimated in XY males). We find that within-pair functional divergence tracks regulatory and structural divergence between the X and Y members, reflecting both upstream and downstream mechanisms that may impact their CFs. We further dissect within-pair functional divergence by ranking all non-gametolog genes by their relative co-expression (i.e., asymmetric coupling) with the X vs. Y member (for each pair and tissue) (Figures 1B, S1). Patterns of asymmetric coupling vary more between gametolog pairs than between tissues, and gene sets associated with specific patterns of asymmetric coupling (across pairs and tissues) are enriched for diverse biological processes, including genome regulation, immune function, and neuronal communication. Moreover, a gene’s cumulative asymmetric coupling to X vs. Y gametologs is associated with sex differences in its expression, co-expression, and capacity to modify risk for sex-biased disease. Taken together, our findings suggest that some gametolog gene pairs do show meaningful functional divergence between their X vs. Y members that is relevant for sex-biased biology.

## Results

### Gametologs exhibit unusually high functional divergence between males and females due to functional divergence between X and Y members

Sex differences in gametolog dosage provide an opportunity to test for functional differences between the X and Y gametologs. In particular, females can express two X members of each pair, whereas males can express one X and one Y member. If X and Y gametologs function similarly at the transcriptomic level, then the combined expression of both members (for each pair) should produce similar CFs in males and females. In contrast, functional divergence between the X and Y members should lead to divergent CFs between males and females. Our analyses found strong support for the latter of these two hypotheses.

For each of 40 tissues, we computed between-sex functional divergence for each gametolog pair using the combined expression of both members in each sex (aCFD_XXF–XYM_) and compared these values to a resampling-based null distribution (see Methods; Figure 1C; see Table S1 for metadata; Table S2). We found that gametolog genes (as a group) exhibit greater average between-sex functional divergence than non-gametolog genes (elevated aCFD_XXF–XYM_ values vs. the null in 37/40 tissues: p_adj_ < 0.05; Figure 2A green points; Table S3). Collectively, gametologs showed the highest levels of functional divergence (aCFD_XXF–XYM_) in digestive tissues (e.g., stomach) and the least in brain tissues (Figure 2A; Tables S2, S3). Certain gametolog pairs showed consistently high aCFD_XXF–XYM_ values across multiple tissue sets, including: *KDM5C/D* and *TXLNG/Y* (tissue Group A; e.g., stomach, transverse colon), *TMSB4X/Y* (Group B; e.g., arteries, thyroid), *USP9X/Y* (Group C; e.g., brain, ileum), and *EIF1AX/Y* (Group D; e.g., heart, liver) (Figures 2B, S2). We illustrate this heterogeneity across gametolog pairs and tissues (pair–tissue) combinations in Figures 2B and 2C. Statistically significant between-sex functional divergence (aCFD_XXF–XYM_, p_adj_ < 0.05) was observed for 108/445 (24%) of all testable pair–tissue combinations (including >=1 tissue for all gametolog pairs and >=1 pair for 36/40 tissues) (Table S2). These findings suggest that gametologs are functionally divergent in males and females.

**Figure 2.**
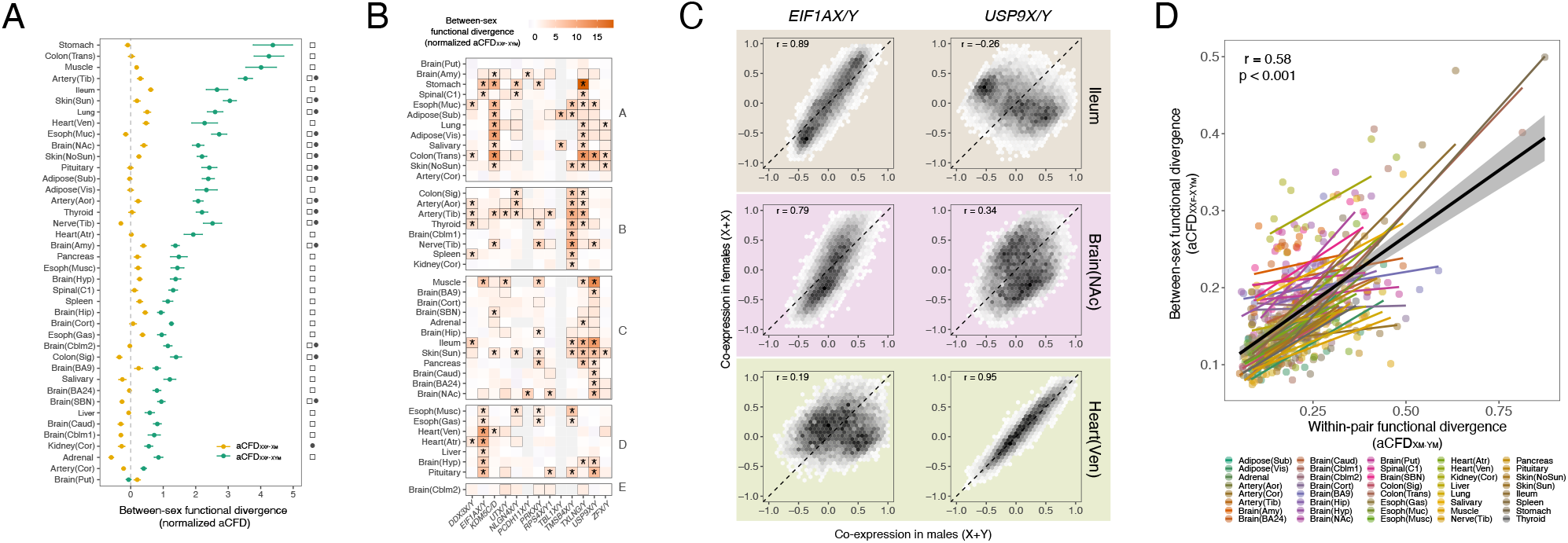
Gametolog gene pairs show prominent functional divergence between males and females. a. Mean (+/-standard error) between-sex functional divergence measures per tissue (averaged across gametolog pairs), estimated by summing X and Y gametolog expression in males (normalized aCFD_XXF–XYM_ in green, per gametolog values shown in Figure 2B) or ignoring Y member expression in males (normalized aCFD_XXF–XM_ in yellow). Open squares indicate tissues where gametologs exhibit higher mean normalized aCFD_XXF–XYM_ values compared to a null distribution (p_adj_ < 0.05, see Methods). Filled circles indicate tissues where normalized aCFD_XXF–XYM_ values are significantly higher than normalized aCFD_XXF–XM_ values (paired t-tests: p_adj_ < 0.05, see Methods). Mean normalized aCFD_XXF–XYM_ across tissues: gametologs = 1.75 [-1.51-18.92], non-gametologs: 0 [-2.75-8.59]. b. Patterning of between-sex functional divergence (normalized aCFD_XXF–XYM_) across gametolog pairs and tissues. Tissues are ordered according to hierarchical clustering of row values and tissue groups A-E were derived from the resulting dendrogram (see Methods). Tiles with black outlines indicate nominal p < 0.05 (vs. co-expression-matched nulls, see Methods). Asterisks indicate p_adj_ < 0.05 (vs. co-expression-matched nulls, see Methods). c. Illustrative examples of CFD for selected tissues and gametolog pairs. Hex-plots show distribution of genes based on their relative co-expression with total expression of the XX gametolog pair in females and the XY gametolog pair in males. Dashed diagonal lines show the identity line (i.e., identical co-expression with the gametolog pair across sexes). Results are shown for two different gametolog pairs (*EIF1AX/Y* and *USP9X/Y*) in three different tissues (ileum, nucleus accumbens, and heart ventricular muscle) (colors correspond to the legend in Figure 2D). Note how *EIF1AX/Y* (left column) shows a very similar co-expression profile with other genes in males and females for the ileum and brain (cross-gene r >= 0.8), but not for the heart ventricles (cross-gene r = 0.2), whereas the opposite is seen for *USP9X/Y* (right column; heart ventricles: cross-gene r > 0.9; ileum and brain: cross-gene r < 0.4). d. Gametolog pairs with greater functional divergence between XX females and XY males (i.e. greater aCFD_XXF–XYM_; y-axis) show greater functional divergence between their X and Y members in males (i.e. greater aCFD_XM–YM_; x-axis). Regression lines are provided for each tissue (see legend) and across all data points (black line, confidence interval is shaded).

Strikingly, gametologs no longer showed elevated between-sex functional divergence when expression of the Y member was dropped from the CF calculation in males. Specifically, when comparing CFs for just the X member(s) of each gametolog pair (using aCFD_XXF–XM_) (Methods; Figure 1C), the average functional divergence of gametolog genes became statistically indistinguishable (p_adj_ > 0.05) from the null gene sets for all 40 tissues (Figure 2A yellow points; Table S3). This finding suggests that gametolog CFs are unusually dissimilar between males and females (i.e., when using aCFD_XXF–XYM_) because female CFs are derived solely from X members while male CFs reflect a combination of dissimilar CFs for X vs. Y members in males. To directly test this hypothesis, we: i) used within-male analyses to compute functional divergence between the X and Y members of each gametolog pair in each tissue (i.e., within-pair CFD, aCFD_XM–YM_) (Figure 1C); and ii) asked whether variation in this metric (aCFD_XM–YM_) predicts the degree of functional divergence between sexes (aCFD_XXF–XYM_) across pairs and tissues. We observed a significant positive correlation between these two metrics (r = 0.58, p < 2.2e-16) (Figure 2D; Tables S2, S4), suggesting that gametolog functional divergence between the sexes is driven by functional divergence between the X and Y gametologs. Sensitivity analyses established that this correlation holds when allowing for tissue-specific effects and estimating CFD in a signed manner see Methods; Tables S2, S4; Figure S3).

As found for between-sex functional divergence (Figure 2B), the degree of functional divergence between X and Y members varied across pairs and tissues (Figures 3A, S2; Table S2). Some pairs showed significantly elevated within-pair functional divergence (aCFD_XM–YM_) in many tissues (e.g., *TXLNG/Y* in 25/43 tissues), while others displayed more tissue-specific elevations (e.g., *PCDH11X/Y* in 2/16 tissues; *ZFX/Y* in 3/43) (Figure 3A, Table S2). We also observed that some tissues show high aCFD_XM–YM_ across multiple gametolog pairs (e.g. adipose: 9/12 pairs; colon: 8/11; stomach: 8/10), while others only showed significantly elevated values for specific gametolog pairs (e.g., amygdala: *PRKX/Y* only; substantia nigra: *USP9X/Y* only; liver: *KDM5C/D only*) (Figure 3A, Table S2). However, overall patterns of functional divergence differed more across pairs than across tissues (ANOVA model: pair: F = 11.98, p < 2.2e-16; tissue: F = 2.09, p = 0.0001), suggesting that within-pair functional divergence largely reflects tissue-wide, sequence-based regulatory and coding dissimilarities between the X and Y members.

**Figure 3.**
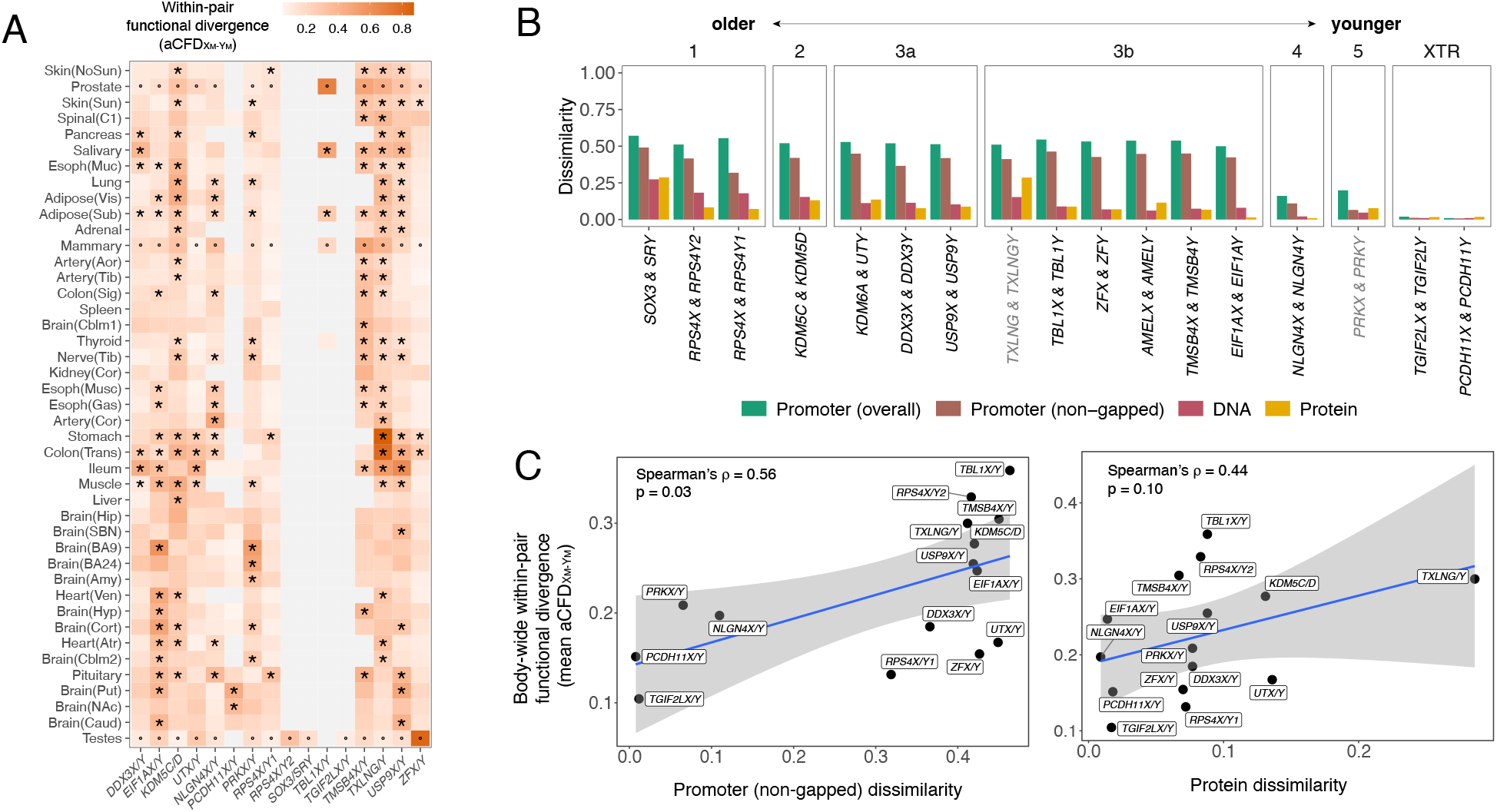
Within pair functional divergence is predicted by regulatory and structural divergence between the X and Y members. a. Absolute within-pair functional divergence between the X and Y members of each gametolog pair per tissue (aCFD_XM–YM_) as calculated in males. Tissues are ordered according to hierarchical clustering. Asterisks indicate instances where the 95% confidence intervals do not overlap between absolute within-pair and between-sex (X member-only) functional divergence (i.e., aCFD_XM–YM_ vs. aCFD_XXF–XM_) and the former is greater (following a resampling approach, see Methods). Small circles indicate that a comparison to between-sex functional divergence was not available because the tissue is sex-specific or sex-differentiated. b. Promoter (overall and non-gapped), DNA, and protein dissimilarity measures for each gametolog pair (see legend). Evolutionary strata from [91] are indicated at the top of the figure. Pairs in gray are those with pseudogenized Y-members. c. Mean absolute within-pair functional divergence (mean aCFD_XM–YM_) vs. promoter divergence (non-gapped) (left) and protein divergence (right). Each point represents one gametolog pair. The regression line (blue line) and its confidence intervals (shaded area) are provided.

### Functional divergence within gametolog pairs is predicted by regulatory and structural divergence over evolutionary time

The analyses above used co-expression fingerprint (CF) analysis to reveal that gametologs collectively show higher functional divergence between males and females than other genes (Figures 2A, 2B, 2C), and that this partly reflects functional divergence between the X and Y members (Fig 2D). The fact that some gametolog pairs tend to show higher within-pair divergence (aCFD_XM–YM_) across multiple tissues (Figure 3A) motivated us to next probe genomic properties that could explain why pairs vary in their divergence. Specifically, we reasoned that higher within-pair functional divergence could reflect: i) sequence divergence in upstream regulatory regions, subjecting X and Y members to different patterns of co-regulation with other genes; and/or ii) sequence divergence in coding regions and the resulting proteins, leading X and Y members to have more divergent downstream effects on the expression of other genes. To test these hypotheses, we scored each gametolog pair for dissimilarity between its X and Y members in terms of their promoter, coding, and protein sequences (Methods). We profiled these features across gametolog pairs (Figure 3B; Table S5) and compared them to cross-tissue (“body-wide”) estimates of within-pair functional divergence (i.e., average aCFD_XM–YM_ across tissues for each pair) (Figure 3C).

As previously reported [7,9,14,19], we found that the X and Y members of gametolog pairs that stopped recombining further back in evolutionary time show more divergent coding and protein sequences, relative to members of their “younger” counterparts (Figure 3B). We also discovered that the same phenomenon holds for promoter sequence dissimilarity (Figure 3B). Although regulatory and structural dissimilarity measures tended to be correlated with each other across pairs (Figure S4; Table S6), pairs with greater body-wide functional divergence (mean aCFD_XM–YM_) also exhibited more dissimilar promoter regions (non-gapped) (*ρ* = 0.56, p = 0.03) and protein sequences (*ρ* = 0.44 p = 0.10) (Table S5). High aCFD_XM–YM_ and protein divergence values for *TXLNG/Y* (Figures 3A, 3C) are consistent with the pseudogenization of *TXLGNY*. Although *PRKY* is also predicted to be a pseudogene, this pair does not exhibit remarkably high aCFD_XM–YM_ or protein divergence values (Figures 3A, 3C), which may reflect recent gene conversion in this area [45]. These results suggest that greater functional divergence between the X and Y members of some gametolog pairs primarily reflects more divergent regulatory mechanisms.

### Delineating the polarity and biological patterning of functional divergence between X and Y gametologs

Thus far, we quantified within-pair functional divergence using the absolute difference between the co-expression fingerprints of their X and Y members (aCFD_XM–YM_) (Figure 1C). This unipolar index (low → high divergence) represents overall functional divergence for each gametolog pair. However, this measure collapses two important axes of information which we next sought to characterize: i) whether global co-expression with all other genes is stronger for the X or the Y member of each gametolog pair (estimated using the signed CFD for each gametolog pair (sCFD_XM–YM_) – henceforth “signed divergence”) (Figure 1C); and ii) which genes are more highly co-expressed with the X vs. Y member of each gametolog pair in each tissue (provided by the full vector of signed gene-level measures of differential co-expression with each X vs. Y member – henceforth “asymmetric coupling”) (Figure 1C).

We first computed signed divergence estimates (sCFD_XM–YM_) (Figure 1C) for each of N=483 gametolog pair–tissue combinations and verified that these are complementary to and non-redundant with absolute within-pair divergence scores (aCFD_XM–YM_) (Figure 4A). For example, pair–tissue combinations with similarly large absolute divergence scores (aCFD_XM–YM_) can show opposing directions of signed divergence (sCFD_XM–YM_) (Figure 4A), indicating a stronger transcriptome-wide co-expression to the X or Y member (Figure 4A). There are also pair–tissue combinations with large absolute divergence scores but near-zero signed divergence scores (Figure 4A), indicating a counterbalancing between genes that are more strongly co-expressed with the X or Y member. The complementarity of absolute and signed divergence scores was also demonstrated by their differential relationships with previously well characterized aspects of gametolog expression, including within-pair (X vs. Y) co-expression and relative expression (Figures S5A-I) [20].

**Figure 4.**
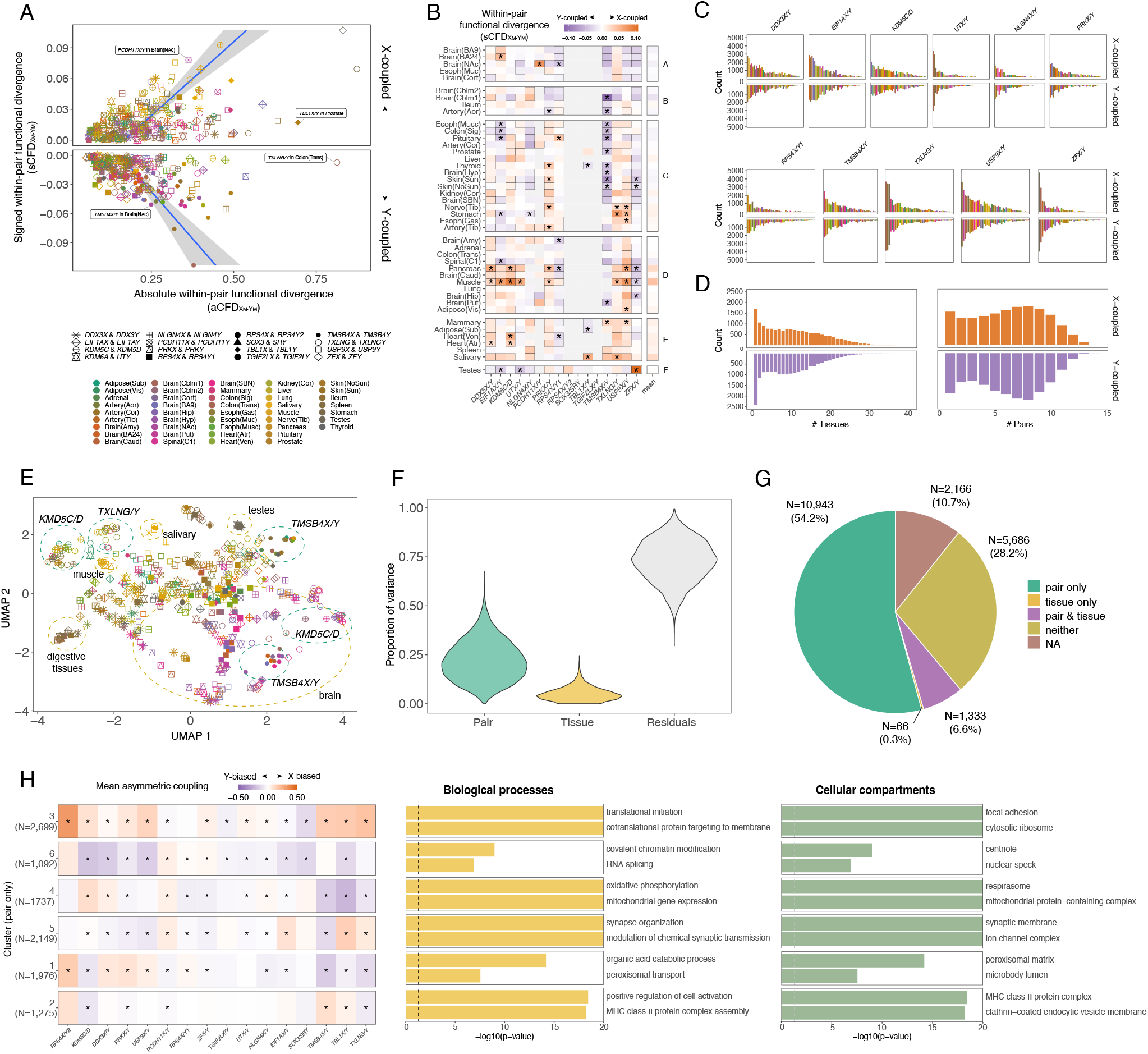
Genes showing asymmetric coupling to X vs. Y gametologs are numerous and enriched within specific biological pathways. a. Signed within-pair functional divergence (sCFD_XM–YM_) as a function of absolute within-pair functional divergence (aCFD_XM–YM_). Colors represent tissues and shapes represent gametolog pairs (see legend). Regression lines are provided for pair–tissue combinations with negative or positive sCFD_XM–YM_ values. Examples are noted for pair–tissue combinations with: i) high aCFD_XM–YM_ but low sCFD_XM–YM_ (*TBL1X/Y* in prostate; *TXLNG/Y* in colon); or ii) similar aCFD_XM–YM_ but divergent sCFD_XM–YM_ (*PCDH11X/Y* vs. *TMSB4X/Y* in nucleus accumbens). b. Signed within-pair functional divergence (sCFD_XM–YM_) for each pair and tissue. Tissues are ordered according to hierarchical clustering and tissue groups A-E were derived from the resulting dendrogram (see Methods). Tiles with black outlines indicate nominal p < 0.05 (estimate = 0; following resampling approach, see Methods). Asterisks indicate p_adj_ < 0.05 (estimate ≠ 0; following resampling approach, see Methods). Mean value column (to the right) is included for visualization purposes only. Note that brain tissues tended to show greater mean Y-coupling, while other tissues showed greater mean X-coupling. c. Count of genes with significant asymmetric coupling (CLIP p_adj_ < 0.05; subsampled to an equal same sample size across tissues, see Methods; minimum male N = 66 in kidney, Figure 1A) per pair and tissue. Only shown for broadly expressed pairs (i.e., those expressed in >50% tissues). Colors indicate tissue (see Figure 4A legend). d. Left: count of genes with significant asymmetric coupling (CLIP p_adj_ < 0.05; subsampled to N = 66 males) in at least one pair across N tissues. Color indicates direction of coupling (see Figure 4B legend). Right: count of genes with significant asymmetric coupling (CLIP p_adj_ < 0.05; subsampled to N = 66 males) in at least one tissue across N pairs. Color indicates direction of coupling (see Figure 4B legend). e. UMAP plot of asymmetric coupling values. Each point is one pair–tissue combination (see Figure 4A legend). Dashed circles are colored according to Figure 4F. Note that pairs tend to cluster together, although there is some clustering across pairs within certain tissues. f. Violin plots depicting the proportion of variance explained (for N = 7,919 genes with asymmetric coupling values for all pair–tissue combinations) by each factor (from variance partitioning analysis; see Methods). Note that a higher proportion of variance in asymmetric coupling values is explained by gametolog pair than by tissue (mean variance explained by pair = 22%; tissue = 5%). g. Pie chart depicting the proportion of expressed genes whose asymmetric coupling values (across pair–tissue combinations) are predicted by pair, tissue, both, or neither (from ANOVA; see Methods). “NA” indicates genes that were not included since they are only expressed in 1 tissue. Note that for most genes, variation in asymmetric coupling values is predicted by gametolog pair alone (for N = 10,943, 54.2% genes expressed in any tissue). h. Left: Mean asymmetric coupling values with each gametolog pair for genes in each cluster (averaged across tissues). N=6 clusters were derived from N=10,943 genes whose asymmetric coupling values were predicted by gametolog pair only (see Figure 4G) (two clusters with < 10 genes were removed). Clusters and pairs are ordered according to hierarchical clustering. Asterisks indicate p_adj_<0.05 (vs. null, see Methods). Middle/right: Top 2 enriched biological processes and cellular compartments for each cluster.

Signed divergence scores showed marked variation across pair–tissue combinations and differed more across pairs than across tissues (ANOVA model: pair: F = 11.40, p < 2.2e-16; tissue: F = 2.89, p = 3.24e-08; Figure 4B). A resampling approach (Methods) revealed statistically significant (p_adj_ < 0.05) signed divergence scores for 63/483 (13%) of all pair–tissue combinations (N = 32 X-biased; N = 31 Y-biased) (including >=1 tissue for all gametolog pairs and >=1 pair for 29/43 tissues) (Figure 4B). The testes showed a distinct pattern of signed divergence (sCFD_XM–YM_) according to hierarchical clustering, driven in part by a distinct pattern of genome-wide coupling to *ZFX* vs. *ZFY* (Figures 4B, S2; Table S2).

We next sought to identify genes that are differentially co-expressed with X vs. Y members of gametolog pairs (and their associated biological processes). To achieve this, we ranked all expressed genes by their contribution to the signed divergence value for each pair–tissue combination (Figure 1C). This produced N = 483 ranked vectors of the directional (X minus Y) Δ co-expression (“asymmetric coupling”) values for each expressed gene, which indicate whether a gene is more strongly co-expressed with the X member (positive value) or Y member (negative value) of the gametolog pair in question (Figure 1; Table S7). Gametolog pairs and tissues varied in the number of genes showing statistically significant asymmetric coupling (identified using an established “correlation by individual level product” (CLIP) approach [40]) (Methods; Figure 1; Figure 4C; Tables S7-10; Figures S6E-G), with the most differentially coupled genes detected in the digestive tissues (averaged across pairs) and – among broadly expressed pairs – *USP9X/Y* (averaged across tissues) (Figure 4C; Tables S7-10). Almost all expressed genes exhibited significant asymmetric coupling with at least one pair–tissue combination (N = 18,902/20,195 [94%]; N = 18,296/18,902 >=1 Y-biased coupling; N = 16,904/18,902 >=1 X-biased coupling) (Tables S7-10), and X chromosome genes were more often asymmetrically coupled than autosomal or Y chromosome genes (chi-squared test: p < 2.2e-16) (Figure S7). Comparisons of asymmetrically coupled genes across pairs and tissues (Figures 4D, S6A-D) suggest that genes tend to be coupled to a few gametolog pairs consistently across multiple tissues.

Three complementary tests established that transcriptome-wide signatures of asymmetric coupling tend to differ more between gametolog pairs than between tissues: dimensionality reduction (Figures 4E, S8), variance partitioning (Figure 4F; Table S12), and ANOVA (Figure 4G; Table S13). This pattern held even when similar tissues were collapsed (see Methods; Figure S8D-F; Table S12). In other words, genes tend to show similar patterns of asymmetric coupling with a given gametolog pair across multiple tissues. However, there were some notable exceptions to this rule, whereby highly similar patterns of asymmetric coupling were observed across multiple gametolog pairs within certain tissues (e.g., the salivary glands, testes, and muscle tissue), suggestive of tissue-specific regulatory processes (Figures 4B, 4E, S8, S9).

To characterize biological processes associated with asymmetric coupling to X vs. Y gametologs, we focused on genes whose asymmetric coupling values were significantly predicted by gametolog pair alone (representing the majority of expressed genes: N = 10,943, 54.2%; Figure 4G; Table S13) (results for other gene sets are in Figure S9). Clustering these genes (based on the similarity of their vectors of asymmetric coupling values) identified 6 distinct gene clusters (excluding two clusters containing <10 genes) (Methods). Each of these clusters exhibited a distinct profile of asymmetric coupling across gametolog pairs (averaged across tissues) (Figures 4H, S9). For example, the largest cluster (Cluster 3; N = 2,699 genes) showed significant X-biased coupling for 11/16 pairs (strongest for *RPS4X/Y2*) and Y-biased coupling for 4/16 pairs (strongest for *SOX3/SRY*), whereas cluster 4 (N = 1,737 genes) showed distinct a distinct profile involving X-biased coupling for 4/16 pairs (strongest for *KDM5C/D*) and Y-biased coupling for 8/16 pairs (strongest for *TBL1X/Y*) (Figures 4H, S9). Gene ontology analysis indicated that these gene clusters exhibit distinct functional annotations. Several clusters were involved in regulatory functions that have already been associated with gametolog genes, including translation (Cluster 3; relevant pairs: *EIF1AX/Y, DDX3X/Y, RPS4X/Y* ; core genes: *EIF1, RPS16, RPL23*), chromatin modification (Cluster 6; relevant pairs: *KDM5C/D, UTX/Y, TBL1X/Y* ; core genes: *ING4, KAT2A, ING5*), and alternative splicing (Cluster 6; relevant pair: *DDX3X/Y* ; core genes: *SNRNP70, RBM6, ZNF638*) [14] (Figure 4H; Tables S14, S15; Figure S9). We also identified a cluster involved in synapse organization that was characterized by greater Y-coupling for most pairs (Cluster 5; core genes: *SYT4, RAB29B, GABRG2*) and an immune-related cluster characterized by greater coupling to *TMSB4X* (vs. *TMSB4Y*) which is expressed in macrophages [46] (Cluster 2; core genes: *APOBEC3G, PLA2G4A, HLA-F*) (Figure 4H; Tables S14, S15; Figure S9).

Collectively, these analyses indicate that genes vary in terms of their relative co-expression to the X or Y gametologs, and that these patterns are shared to some extent across tissues. Genes with similar patterns of asymmetric coupling are associated with: i) the regulatory functions of gametolog genes; and ii) diverse biological processes, including several that are implicated in sex-biased disorders of neurodevelopment and autoimmunity.

### Asymmetric coupling to the X vs. Y gametologs (in males) is associated with sex differences in gene expression and co-expression

The above analyses – conducted within XY males – indicate that genes vary in their profiles of asymmetric coupling with X vs. Y gametologs across pairs and tissues. If the magnitude of asymmetric coupling reflects a gene’s relative co-regulation with and/or its regulation by X vs. Y gametolog genes, then normative sex differences in expression and co-expression of autosomal genes may be partly organized by their asymmetric coupling to gametologs. This suggests that – although there are numerous other factors that jointly regulate sex differences in expression (e.g., sex-biased hormones, transcription factor targeting, and genetic regulators of expression (eQTLs) [47–49]) – there may be detectable proportion of sex-related variation that is attributable to gametolog functional divergence.

To address this idea, we tested whether autosomal genes that tend to be more highly expressed and co-expressed in females vs. males, respectively, show greater overall coupling to the X vs. Y gametologs, respectively. First, for each autosomal gene in each tissue, we estimated the magnitude of sex differences in expression (“sex-biased expression”) (Methods; Table S20) and compared these values to summary measures of asymmetric coupling (“overall asymmetric coupling”). The latter represent weighted average asymmetric coupling values, which account for each gene’s correlations with and the expression levels of each X and Y gametolog (Methods; Figure S1). Across tissues, we found that: i) autosomal genes with significantly female-vs. male-biased expression levels (local false sign rate (LFSR) < 0.05; see Methods) tended to exhibit asymmetric coupling (summed CLIP: p_adj_ < 0.05) to X vs. Y gametologs, respectively (65% concordant; Fisher’s exact test: OR = 3.61, p < 0.01); and ii) these measures were positively correlated across genes (*ρ* = 0.34; p < 0.01) (Figure 5A). This relationship was also present within most tissues (Table S21; Figure S10). We then compared sex-biased *co-expression* levels to tissue-specific overall asymmetric coupling values. The latter represent overall connectivity [50] and were obtained from a previous analysis of the GTEx data [51]. We detected a pattern similar to that observed for sex-biased expression: across N=22 tissues examined in [51], genes with X-biased coupling tended to show female-biased (but not male-biased) co-expression (X-biased + female-biased: OR = 1.15, p = 0.02; X-biased + male-biased: OR = 0.98, p = 0.67) while Y-biased coupling was weakly associated with male-biased (but not female-biased) co-expression (Y-biased + male-biased: OR = 1.03, p = 0.35; Y-biased + female-biased: OR = 0.87, p = 0.98) (Figure 5B). This pattern was also detected within certain tissues (Table S22). Furthermore, we detected a complementary pattern linking asymmetric coupling to sex differences in co-expression: autosomal gene sets defined by contrasting directions of asymmetric coupling (i.e., those with greater overall coupling to X gametologs vs. those with greater overall coupling to Y gametologs) were more weakly co-expressed with each other in XY males (where both X and Y gametolog influences are operative) compared to XX females (Methods) (paired t-test of mean co-expression [scaled within each sex]: p = 3.43e-11; mean difference = 0.14 [XX > XY]) (Figure 5C). Taken together, these results suggest that asymmetric coupling to the X vs. Y gametologs, combined with sex differences in sex chromosome dosage, contribute to sex differences in gene expression and co-expression across the human body.

**Figure 5.**
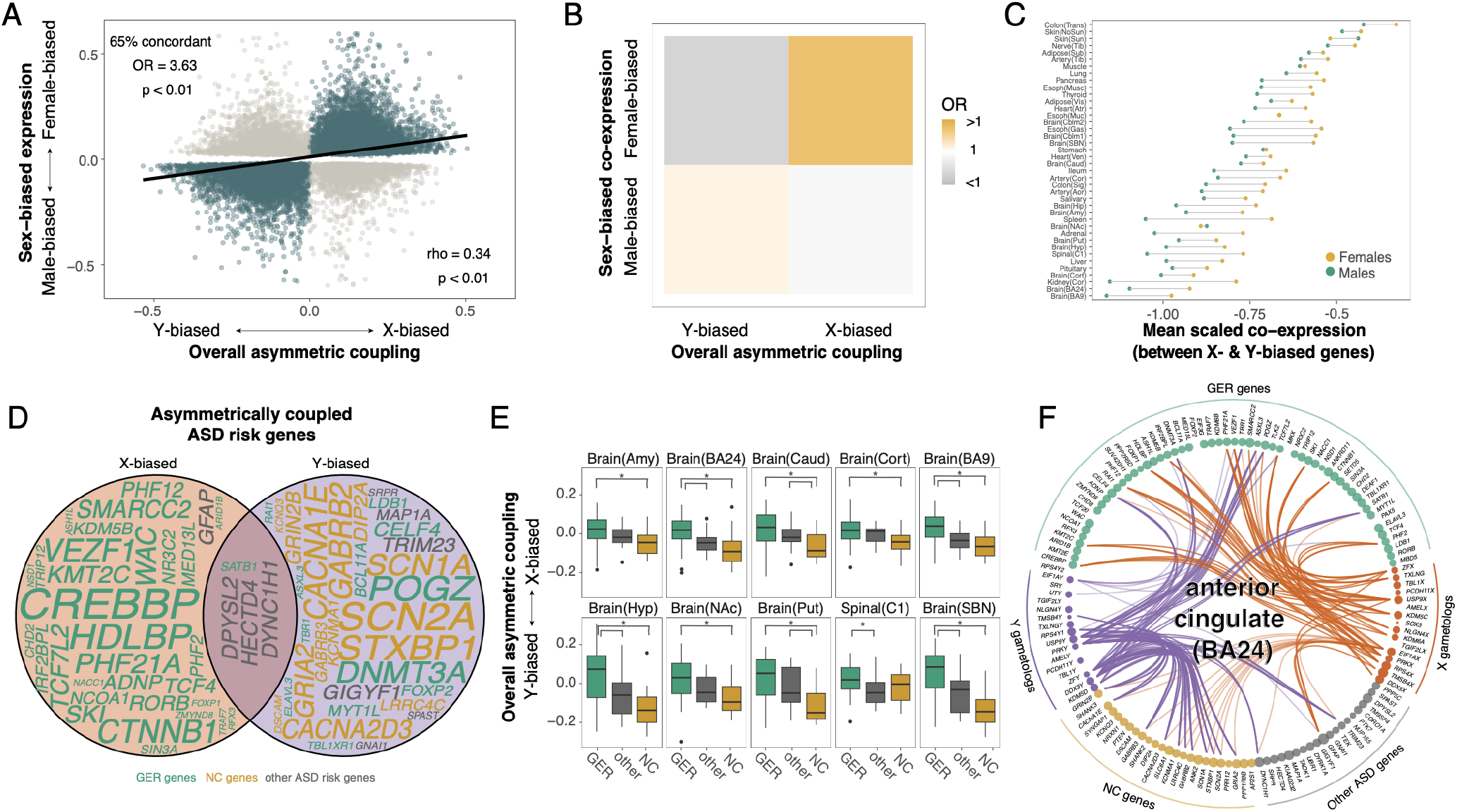
Asymmetric coupling to the gametologs is related to sex differences in gene expression, co-expression, and genetic contributions to ASD. a. Sex-biased gene expression (y-axis; positive = female-biased, negative = male-biased; values = mashr *β*) as a function of asymmetric coupling (x-axis; positive = X-biased, negative = Y-biased). Each point represents the values for one gene in one tissue. Only genes with significant sex-bias (LFSR < 0.05) and overall asymmetric coupling (overall CLIP p_adj_ < 0.05) are included (N=23,053 observations across tissues; N=7,690 unique genes). Regression line is provided. Colors indicate genes with sex-bias and asymmetric coupling patterns in the expected (“concordant”) direction (in green) or the opposite (in gray). b. Enrichment results between overall asymmetric coupling and sex-biased co-expression (see Methods). OR = odds ratio. c. Mean scaled co-expression values between all pairs of X-biased genes and Y-biased genes in each tissue in males (green) versus females (yellow). d. Venn diagram of ASD risk genes that are significantly asymmetrically coupled (overall CLIP p_adj_ < 0.05) in at least one brain tissue. Gene names are colored according to their functional category (from [58]; see legend below plot; GER = gene expression regulation, NC = neuronal communication). N = 32 genes were only ever X-coupled (in >=1 tissue; N = 31 GER, N = 1 other); N = 32 genes were only ever Y-coupled (in >=1 tissue; N = 14 NC, N = 12 GER, N = 6 other); N = 4 genes showed opposing coupling patterns across brain tissues (N = 1 GER; N = 3 other). Size of gene names is proportional to the number of brain regions in which the gene shows significant asymmetric coupling (tissue-specific overall CLIP p_adj_ < 0.05). Overall chi-squared test: p = 3.45e-07; post-hoc Fisher’s exact tests p_adj_ < 0.05 for X-biased vs. Y-biased, X-biased vs. neither/both (unbiased), GER vs. NC, GER vs. other. e. Boxplots of asymmetric coupling values for ASD risk genes in different functional groups (regardless of CLIP p_adj_ value) for N = 10 brain tissues. Colors correspond to those in Figure 5D. f. Example of divergent asymmetric coupling across N = 101 ASD risk genes in the anterior cingulate cortex (BA24). Each dot represents an ASD risk gene (grouped by functional category; colors correspond to those in Figures 5D and 5E) or a gametolog gene (separated by chromosome; colors correspond to those in Figures 4B and 4D). Dot size is proportional to the mean adjusted expression level of that gene in BA24. For each ASD risk gene with significant overall asymmetric coupling in BA24 (overall CLIP p_adj_ < 0.05; N = 32 genes), a connection is depicted with either the X or Y member of each gametolog pair to which the ASD risk gene is asymmetrically coupled (CLIP p_adj_ < 0.05) (i.e., if an ASD risk gene is asymmetrically coupled to *ZFX/Y* and more strongly co-expressed with *ZFX* vs. *ZFY*, then this gene’s connection to ZFX is depicted here; colors of the connecting lines correspond to the X (orange) or Y (purple) gametologs). No connections are shown for ASD risk genes with non-significant overall asymmetric coupling (overall CLIP p_adj_ < 0.05) or between ASD risk genes and individual gametolog pairs to which they are not asymmetrically coupled (CLIP p_adj_ < 0.05). The opacity of each connecting line reflects the overall asymmetric coupling pattern of the given ASD risk gene (e.g., *PHF21A* (towards the top of the plot) exhibits overall X-biased coupling in BA24, so its N = 4 connections to X gametologs (*PRKX, RPS4X, USP9X, ZFX*) are shaded darker than its N = 1 connection to a Y gametolog (*EIF1AY*); *TBR1* (also towards the top of the plot) exhibits overall Y-biased coupling in BA24, so its N = 4 connections to Y gametologs (*DDX3Y, PRKY, USP9Y, ZFY*) are shaded darker than its N = 1 connection to an X gametolog (*EIF1AX*)).

### Asymmetric coupling to X vs. Y gametologs intersects with genetic risk for a sex-biased disorder

The relationships between asymmetric coupling and normative sex differences in gene expression and co-expression (Figures 5A, 5B) suggest a potential role for asymmetric coupling in shaping genetic risk for sex-biased diseases. This reasoning hinges on observations that: i) disease genes tend to be more highly expressed and occupy more central positions in co-expression networks [52–55]; and ii) genes exhibiting sex-biased expression and co-expression overlap with genetic risk for sex-biased diseases in a directional manner [51,56]. As an initial test of this idea, we focused on autism spectrum disorder (ASD) since: i) ASD is a paradigmatically male-biased condition [57]; ii) there are well-established sets of high-impact ASD risk genes [58]; iii) the GTEx provides multiple brain tissues for characterizing the asymmetric coupling of these genes (Figure 1A); and iv) there is emerging evidence that sex-biases in penetrance may vary across functional groups of ASD risk genes [59]. High-confidence ASD risk genes (implicated by rare variants) fall into two major subclasses that differ in their biological functions, cellular localizations, and spatiotemporal expression patterns in the brain [58,60–62]: nuclear-expressed genes involved in gene expression regulation (GER) and synaptically expressed genes involved in neuronal communication (NC). We therefore tested these two subsets of ASD risk genes for asymmetric coupling with gametologs.

Our analyses considered N = 102 ASD risk genes identified by Satterstrom and colleagues [58], 80% of which are involved in GER (N = 58) or NC (N = 24) (Table S23). The remaining 20% (“other” risk genes; N = 20) are associated with e.g., the cytoskeleton (Table S23). We detected almost all ASD risk genes (101/102) in at least one GTEx brain tissue (N = 13 total). We first classified these genes into three categories based on their patterns of overall asymmetric coupling (overall CLIP p_adj_ < 0.05) across brain regions, including those that: i) are only ever X-biased (in >=1 region; N = 32; “X-biased ASD genes”); ii) are only ever Y-biased (in >=1 region; N = 32; “Y-biased ASD genes”); or iii) show a regionally mixed pattern (N = 4) or exhibit no bias (N = 33) (“unbiased ASD genes”). We found that these categories were associated with ASD risk gene functional categories (chi-squared test: p = 3.45e-07): X-biased ASD genes tended to be GER-related (31/32 = GER, 1/32 = other), Y-biased ASD genes tended to be NC-related (14/32 = NC, 12/32 = GER, 6/32 = other), and unbiased ASD genes were involved in all functional categories (14/37 = GER, 13/37 = other, 10/37 = NC) (Figure 5D; Table S23). In a complimentary test, we established that GER-related ASD risk genes exhibited greater X-biased coupling on average, while NC ASD risk genes exhibited greater Y-biased coupling (regardless of the significance of their asymmetric coupling) (ANOVA p_adj_ < 0.05 in 10/13 tissues; Tukey’s HSD p_adj_ < 0.05: GER > NC in 9/13, GER > other in 5/13; other > NC in 2/13) (average of mean asymmetric coupling values across tissues: GER = 0.03, other = −0.02, NC = −0.05) (Figure 5E; Tables S23, S24). X-biased coupling among GER genes was driven by *USP9X, UTX/KDM6A*, and *EIF1AX* (Table S7; Figure 5F), which indirectly regulate gene expression via translational regulation of other genes [14,63]. Y-biased coupling among NC genes was driven by *USP9Y* and *ZFY* (Table S7; Figure 5F), the latter of which is a known transcription factor that activates gene expression throughout the genome [21].

These findings for ASD open the door for broader consideration of asymmetric coupling with gametologs as a sex-biased aspect of genetic risk across medicine.

## Discussion

Our study harnesses co-expression fingerprint (CF) analysis as a fresh tool to address the long-debated question of functional equivalence between the X and Y gametologs. To date, direct experimental comparisons between X and Y gametologs have been limited to a few pairs and functional read-outs, yielding isolated assertions of functional convergence [21,29] or divergence [25,26,30–33] and conflicting conclusions from different assays of “function” (e.g., *DDX3X* and *DDX3Y* have convergent effects on protein synthesis [29] but divergent effects on RNA metabolism [25]). The computational approach used here provides a necessary accompaniment to these experimental studies by using genome-wide co-expression data to systematically detail the magnitude and nature of gametolog functional divergence across pairs and tissues. This strategy advances current understanding in several key directions.

First, we establish that gametolog genes collectively show high functional divergence between males and females (relative to other genes). We then show that this phenomenon is driven by divergence between the X and Y members (assayed in males), which is greatest among evolutionarily distant pairs with more divergent regulatory sequences. Furthermore, the magnitude of functional divergence – both between the sexes and the X vs. Y members – is highly variable across pairs and (to a lesser extent) across tissues. This variation is consistent with isolated experimental studies (e.g., low aCFD_XM–YM_ for *ZFX/Y* in most tissues mirrors the recent finding that knockdown of *ZFX* or *ZFY* in fibroblasts has similar effects on the expression of other genes [21]). The pair- and tissue-specific patterns of functional divergence described here can be used to guide future experimental tests of gametolog divergence. For example: i) *USP9X/Y* exhibits high functional divergence across brain tissues (Figures 2B, 3A, 4B), suggesting it should be prioritized for analysis of functional divergence in the brain; ii) *EIF1AX/Y* exhibits high functional divergence in the heart (Figures 2B, 3A, 4B), consistent with previous work linking this gametolog pair to sex-biased expression and heart disease [20]; and iii) the testes show a distinct genome-wide coupling bias towards *ZFX* (vs. *ZFY*) (Figures 2B, 3A, 4B), suggesting a mechanism underlying the severe spermatogenic defects caused by mutations to *ZFX* in humans [64].

Second, our computational approach provides unique insights into gametolog functional divergence by providing tissue-specific, gene-level estimates of asymmetric coupling to the X vs. Y gametologs. Importantly, recent experimental work converges with this computational assay: genes differ in their magnitude of expression level change following *ZFX* vs. *ZFY* knockdown [21], implying that individual genes are more tightly co-regulated with and/or regulated by individual X vs. Y gametologs. Our analyses reveal that almost all expressed genes are asymmetrically coupled to the X or Y member of at least one gametolog pair in one tissue, suggesting broad potential impacts of gametolog functional divergence across diverse biological domains. In fact, we found that patterns of asymmetric coupling are associated with specific biological processes, including synaptic function and immune activation, which are known hotspots for sex-biased biology and disease risk [1,65]. Furthermore, we show that sex differences in gene expression and co-expression – which are likely to reflect complex interactions between diverse genetic, endocrine and experiential factors [66] – can be partly predicted from gene-level measures of asymmetric coupling to X vs. Y gametologs (quantified in males). This finding suggests that categorical sex differences in the expression levels of X vs. Y gametologs [13–20] combine with the functional divergences we report herein to constitute an important and previously underappreciated source of sex-biased biology.

Finally – using ASD as an illustrative example – we show that two functionally distinct sets of ASD risk genes show divergent patterns of X-vs. Y-biased coupling in the human brain. These findings nominate asymmetric coupling as an additional mechanism underlying sex differences in genetic risk for ASD, and highlight specific gametologs (e.g.,*USP9X/Y*) for further investigation. Moreover, the approaches demonstrated for ASD risk genes can readily be applied to any other gene sets of interest for sex-biased disease (e.g., genes implicated by genetic association or transcriptomic alteration), opening up new candidate mechanisms for well-reported but poorly understood sex differences in disease prevalence, presentation, and prognosis [1–3].

Our findings should be considered in light of some methodological and conceptual limitations. First, the GTEx resource consists of bulk RNA-Seq data, precluding our ability to characterize gametolog functional divergence within specific cell types. However, given that transcriptome-wide patterns of asymmetric coupling to individual gametolog pairs tend to be similar across tissues composed of diverse cell types (Figures 4E-4G), these patterns may also be similar across tissue-specific cell types. Future work using single cell data will be able to address this. Second, the current study cannot identify which evolutionary mechanisms produced the functional divergences we observe between X and Y gametologs in humans. These non-mutually exclusive mechanisms include positive and relaxed purifying selection, which promote the spread and fixation of beneficial mutations or reduce the removal of deleterious mutations, respectively. Both mechanisms are consistent with reports that the coding sequences of Y gametologs evolved faster than those of X gametologs [6,10,67]. However, they suggest different functional implications for X–Y divergences, evidence of which varies across gametolog pairs and evolutionary timescales. For example: i) positive selection on certain Y gametologs may reflect novel, male-specific functions [68,69]; and/or ii) relaxed purifying selection on some Y (vs. X) gametologs may have lessened their efficacy (leading to evolutionary reductions in Y gametolog expression [6,28], higher combined gametolog expression in XY vs. XX individuals [10,17,20,28], and stronger upregulation of gametologs from increased Y (vs. inactive X) chromosome dosage [21]). Notably, possible reductions in translational efficiency among Y (vs. X) gametologs [20] is unlikely to explain the divergent CFs we observe for X and Y gametologs, as this should result in similar X vs. Y member regulation across XY males. Finally, these data are cross-sectional and limited to adult individuals, so we are unable to address questions regarding the timing of gametolog functional divergence across development and the postnatal lifespan.

Taken together, our findings identify and characterize functional divergence between the X–Y gametologs. This divergence is greatest among evolutionary distant gametolog pairs and manifests as autosomal genes showing coordinated patterns of asymmetric coupling with X vs. Y gametologs. We further show how patterns of asymmetric coupling are enriched for specific functional pathways and gene sets implicated in sex-biased aspects of health and disease, supporting claims that even small differences in X–Y gametolog function and/or regulation may shape phenotypic differences between XX and XY individuals [10,20,21,70]. These insights provide a rich empirical resolution to long-running debates regarding the degree, form, and biological relevance functional divergence between X–Y gametolog pairs.

## Methods

### Read alignment and quantification

We downloaded all GTEx (v8) RNA-Seq BAM files via dbGaP (project #26993) and converted them to FASTQ files using the SamToFastq function from the GTEx RNA-seq pipeline [41]. We filtered out samples that did not pass GTEx QC filters [41], resulting in a dataset of N=17,382 samples spanning N=55 tissues. We then mapped reads to the human transcriptome GRCh38 (Ensembl) using the pseudoaligner kallisto v0.46.2 [71] with sequence-based bias correction (–bias) [20]. Given that sequence homology across the sex chromosomes present in reference genomes/transcriptome can lead to technical mapping errors, we created a modified, sex-specific transcriptome and separately mapped reads from males and females [72]. Specifically, the Y chromosome was removed from the female-specific transcriptome. No alterations were made to the PAR regions since they are not present in the reference Y chromosome. We also confirmed chromosomal sex of individuals by mapping to a non-sex-specific transcriptome and examining Y chromosome gene counts.

We imported the transcript count matrices for males and females into R using the function tximport in the R package tximport [73] and combined them into one count matrix. We summarized transcript counts to the gene level using the appropriate functions in the R package biomaRt and the function summarizeToGene (R package tximport) [73]. This procedure resulted in a 40,321 × 17,382 (p x n) count matrix, where p is the number of genes measured and n is the number of samples (5,798 females; 11,584 males).

The GTEx v8 data includes technical replicates for two tissues: 1) Brain - Frontal Cortex (BA9) and Brain - Cortex; and 2) Brain - Cerebellum and Brain - Cerebellar Hemisphere (GTEx site). We analyzed these replicates separately.

### Filtering and normalization

We first filtered the gene expression matrix to include tissues (vs. blood or cell lines) with N>20 male samples, resulting in 15,418 samples (N=4,794 females, N=10,624 males) across N=43 tissues. Then, we filtered out genes that were very lowly or not detectably expressed by removing any gene with mean CPM<5 (if the gametologs were filtered out here, they were added back to the expression matrix as long as their mean CPM>=1), resulting in a mean of N=12,829 expressed genes per tissue (min=10,228 [skeletal muscle], max=15,946 [testis]; N=20,194 genes expressed in any tissue; Table S1). We then calculated normalization factors using the calcNormFactors function (R package edgeR) and the TMM method, which accounts for differences in sequencing depth, RNA composition, and gene length [74]. Counts were then normalized and converted to log2(cpm) using the voom function (R package limma) [75]. Finally, we identified and removed outlier samples whose mean normalized gene expression correlation with all other samples in the tissue was beyond 3 standard deviations of the mean, resulting in 15,155 samples (N=4,707 females, N=10,448 males; Table S1). For the sex-specific analyses described below, these filtering and normalization steps were also performed within each sex.

### Adjusting gene expression levels for covariates

For each tissue separately and within each sex, we removed the effects of age [AGE] and technical variables (RIN [SMRIN], ischemic time [SMTSISCH], intronic mapping rate [SMNTRNRT]) from the normalized gene expression matrix using the removeBatchEffects function (R package limma) [75]. This function fits a linear model to the expression matrix and then removes the components associated with the covariates, producing an adjusted gene expression matrix. All continuous predictors were scaled and centered prior to linear modeling (mean=0, sd=1).

### Calculating co-expression

To obtain co-expression values within each sex and for each tissue, we estimated Spearman’s rank order correlations of covariate-adjusted expression levels for all pairs of genes using the cor function (R package stats). Given that gene expression correlation matrices exhibit a mean-correlation relationship, in which highly expressed genes appear more correlated than lowly expressed genes [76], we applied spatial quantile normalization (SpQN) to each co-expression matrix to obtain normalized co-expression matrices. This method uses bins to partition the correlation matrix (i.e., genes are sorted by their average expression level) and then applies quantile normalization to each bin. To smooth the normalization, we used a larger bin to approximate the empirical distribution of the inner bin, and applied 1-dimension quantile normalization on the inner bins using its approximated distribution. The target distribution we used here is the bin with the second highest expression level on the diagonal. This was executed using the normalize_correlation function (R package spqn) with the following parameters: ngrp = 20, size_grp = 1000, ref_grp = 18 [76].

To facilitate analyses described below, co-expression vectors in males were estimated using two approaches: 1) co-expression values for the X and Y gametologs were estimated separately; and 2) the co-expression value for each gametolog pair was estimated after adding together the adjusted expression levels of the X and Y gametologs.

### Between-sex co-expression fingerprint divergence

To estimate between-sex CFD, we first removed: i) genes that were not expressed in both sexes; and ii) tissues that are sex-specific or highly differentiated (N=40 tissues retained, excluding testis, prostate, mammary). Then, within each remaining tissue, we: 1) subtracted male and female co-expression matrices, found the absolute value of all values in the resulting matrix, and averaged these values for each gene (aCFD, see Figure 1C); and 2) subtracted male and female co-expression matrices and averaged these signed values for each gene (sCFD, see Figure 1C). For the gametolog genes, this process was repeated using two approaches (the input matrices are described above): i) we estimated between-sex CFD for each gametolog pair using co-expression values derived from adding together the adjusted expression levels of the X and Y gametologs (aCFD_XXF–XYM_); and ii) we estimated sex-dependent CFD for each gametolog pair using co-expression values for the male X gametolog only (aCFD_XXF–XM_).

To test whether gametologs (as a group) tend to have higher aCFD_XXF–XYM_ than non-gametolog genes (results shown in Figure 2A, green), we compared the mean aCFD_XXF–XYM_ values (averaged across the gametologs within each tissue) to a null distribution derived from genes with similar average global co-expression profiles. Within each tissue, we sampled N genes equal to the number of gametolog pairs expressed in that tissue, each within the same decile of mean co-expression to all other genes as the matched gametolog pair. We then estimated the mean aCFD_XXF–XYM_ for these genes and repeated this process 1000 times to obtain a distribution of null mean aCFD_XXF–XYM_ values. In each tissue, the p-value was calculated as the proportion of absolute null mean aCFD_XXF–XYM_ values that were greater than the observed mean aCFD_XXF–XYM_ value for the gametologs. P-values were adjusted across tissues using Benjamini-Hochenberg [77].

To test whether between-sex CFD (as calculated above) was impacted by inclusion of Y gametolog expression levels (in estimating the gametolog CFs in males), we compared mean z-score normalized aCFD_XXF–XYM_ values (including Y-member expression for the gametologs in males) to mean z-score normalized aCFD_XXF–XM_ values (excluding Y member expression for the gametologs in males) within each tissue using paired t-tests (results shown in Figure 2A, yellow). Z-score produces a set of values with a mean = 0 and a standard deviation = 1. P-values were adjusted using Benjamini-Hochenberg [77].

To test whether individual gametolog gene pairs (including Y-member expression for the gametologs in males) exhibit higher between-sex CFD than non-gametolog genes (results shown in Figure 2B), we compared the aCFD_XXF–XYM_ value of each gametolog pair (within each tissue) to a null distribution derived from genes with similar average global co-expression profiles. Specifically, within each tissue and for each gametolog pair, we obtained a null distribution of aCFD_XXF–XYM_ values by sampling 1000 genes within the same decile of mean co-expression (with all other genes) as the gametolog pair. In each case, a p-value was calculated as the proportion of absolute null aCFD_XXF–XYM_ values that were greater than the observed aCFD_XXF–XYM_ value. P-values were adjusted using Benjamini-Hochenberg [77].

Hierarchical clustering of tissues according to normalized aCFD_XXF–XYM_ values (across gametolog pairs) was performed using the pvclust function in the R package pvclust [78]. Groups of tissues (results shown in Figure 2B) were obtained by cutting the resulting dendrogram at height = 0.8. pvclust provides two types of p-values: the Approximately Unbiased p-value (AU; computed by multiscale bootstrap resampling) and the Bootstrap Probability value (BP; computed by normal bootstrap resampling).

### Within-pair co-expression fingerprint divergence

We estimated within-pair CFD using co-expression matrices in XY males. For each gametolog pair–tissue combination, we subtracted the Y-member’s CF from the X-member’s CF, creating a vector which detailed each gene’s “asymmetric coupling” with that gametolog pair in that tissue (see Figure 1C). Co-expression values for each X- and Y-member were Fisher-Z transformed (for normalization and variance-stabilization) prior to subtraction [79] (using the FisherZ function in the R package DescTools) [80]. We then calculated the mean of these values (sCFD_XM–YM_) or of the absolute value of these values (aCFD_XM–YM_) to give pair-level indices of co-expression divergence for each gametolog pair in each tissue.

To test the significance of within-pair sCFD for each pair in each tissue (results shown in Figure 4B), we used the following resampling approach: i) within each tissue, we resampled columns (i.e., individuals) from the male adjusted expression matrix (with replacement) and recalculated the co-expression matrix (applying SpQN normalization, as described above); ii) estimated sCFD_XM–YM_ for each gametolog pair; iii) repeated this process (steps i and ii) 100x (for computational tractability) to create a distribution of sCFD_XM–YM_ values for each pair; and iv) used the sCFD_XM–YM_ distributions for each pair to derive means, standard deviations, confidence intervals, and associated p-values (i.e., whether the mean _sCFDXM–YM_ is significantly different from zero).

To test the significance of within-pair aCFD for each pair in each tissue (results shown in Figure 3A) – we compared the distribution of resampled aCFD_XM–YM_ values to the distribution of resampled aCFD_XXF–XYM_ values (i.e., we compared the distributions of absolute within-pair vs. between-sex CFD). Given that within-pair aCFD values are all positive, we could not apply the method described above for within-pair sCFD (which tests for significant deviations from zero). Specifically, for each pair in each tissue, we: i) applied the resampling procedure described above for signed within-pair CFD (sCFD_XM–YM_) to both aCFD_XM–YM_ and aCFD_XXF–XYM_; and ii) derived confidence intervals (CIs) for both aCFD_XM–YM_ and aCFD_XXF–XYM_ values. Pair–tissue combinations for which the lower CI for aCFD_XM–YM_ was greater than the upper CI for aCFD_XXF–XYM_ were identified as having significantly high within-pair aCFD (aCFD_XM–YM_).

### Identifying genes with significant asymmetric coupling with each gametolog pair in each tissue

The significance of the asymmetric coupling (for each gene to each gametolog pair in each tissue) was estimated using the “correlation by individual level product” (CLIP) method [40], which was developed to identify factors associated with variation in gene expression correlations. To apply this method, we scaled the adjusted expression values (for each expressed gene across all individuals) to a mean of 0 and a standard deviation of 1 (using the scale function in the R base package). Then, for each gametolog pair, the scaled expression level of each (non-gametolog) gene was multiplied by the scaled expression of the X-member and (separately) by the scaled expression of the Y-member within each individual. This produced two vectors for each non-gametolog gene, containing the products of the expression levels of the gene and the X or Y gametolog, respectively, across individuals. To identify genes with significantly greater coupling to the X or Y gametologs, the mean values of these vectors were compared using paired t-tests (using the t.test function in the R package stats) (Figure S1). This procedure was repeated using the summed expression levels for all X gametologs or all Y gametologs to test the significance of the magnitude of overall asymmetric coupling (across all gametolog pairs) for each gene within each tissue (Figure S1). P-values were adjusted within each tissue using Benjamini-Hochenberg [77]. We verified that CLIP t-values were positively correlated with asymmetric coupling values (across genes, per pair–tissue: *ρ* = 0.95, p < 2.2e-16; across genes, expression-weighted average values per tissue; *ρ* = 0.89, p < 2.2e-16).

To compare the number of significantly asymmetrically coupled genes across pairs–tissues, we subsampled each tissue to the minimum number of males (N=66) and recalculated asymmetric coupling values and associated adjusted p-values (using the CLIP method), as described above [40] (results shown in Figure 4C-D). The number of significantly asymmetrically coupled genes was positively correlated across analyses of the total sample and of the subsamples (per tissue: *ρ* = 0.39, p = 0.01; per pair: *ρ* = 0.97, p = 2.77e-10; per pair–tissue: *ρ* = 0.61, p < 2.2e-16) (Figure S6E-G).

### Computing a global asymmetric coupling value across all gametologs for each gene in each tissue

Sex differences in gene expression and co-expression are typically estimated as gene-level properties. To test how these sex differences might relate to gametolog functional divergence (results shown in Figures 5A-C), we sought to create a summary gene-level estimate of asymmetric coupling to all X vs. Y gametologs (within each tissue). To achieve this, we estimated an expression-weighted mean asymmetric coupling value for each expressed gene in each tissue (i.e., “overall asymmetric coupling”), taking into account the expression levels of each X or Y gametolog gene. Specifically, for each expressed gene in each tissue, we: 1) multiplied its correlation with each X gametolog by the adjusted expression level of each X gametolog, summed these products, and divided them by the sum of the adjusted expression level of all X gametologs (to obtain a weighted average correlation of the gene with the X gametologs); 2) repeated this process using the Y gametologs; and 3) subtracted the weighted average correlation of the gene with the Y gametologs from the weighted average correlation of the gene with the X gametologs (Figure S1). Genes were then ranked by these values and these rankings were used in our comparisons to sex-biased gene expression and disease enrichment analyses (see below).

### Estimating structural dissimilarity scores

For each gametolog pair, we estimated promoter sequence, DNA sequence, and protein sequence dissimilarity.

Promoter sequence dissimilarity: We obtained the promoter sequences (2000 bps upstream) for each gametolog using the getSequence function (R package biomaRt) [81] (seqType = ‘gene_flank’; provides the flanking region of the gene excluding the UTRs). For each pair, we aligned these sequences using the pairwiseAlignment function (R package BioStrings), and two measures of promoter dissimilarity were estimated: 1) overall promoter dissimilarity = 1 - the number of matches divided by the length of the alignment (considers gaps); 2) non-gapped promoter dissimilarity = 1 - the number of aligned matches divided by the sum of matches and mismatches (does not consider gaps). While there appeared to be a large shift in promoter dissimilarity between strata 3b and 4, this is likely to reflect a ceiling effect for our measure of promoter dissimilarity.

DNA sequence dissimilarity: For each gametolog pair, DNA divergence scores were obtained from Skaletsky and colleagues [19] and DNA sequence dissimilarity was estimated as the divergence score/100.

Protein sequence dissimilarity: For each gametolog pair, protein divergence scores were obtained from Skaletsky and colleagues [19] and protein sequence dissimilarity was estimated as the divergence score/100.

We first estimated how similar all dissimilarity measures were to each other using Spearman’s rank order correlations (cor function in the R package stats) (Figure S4). Finally, we estimated Spearman’s rank order correlations (using the cor function in the R package stats) between each dissimilarity measure described above and mean absolute within-pair CFD (aCFD_XM–YM_; averaged across tissues for each gametolog pair) (results shown in Figure 3C).

### Deconstructing patterns of asymmetric coupling

We used multiple methods to describe the patterns of asymmetric coupling across genes, including dimensionality reduction (results shown in Figure 4E), variance partitioning (results shown in Figure 4F), and ANOVA (results shown in Figure 4G). These were performed: i) on the whole matrix of asymmetric coupling values (i.e., an N x M matrix; N = # genes; M = # pair–tissue combinations); and ii) on a matrix in which the values for similar tissues were averaged at the organ-level (i.e., “collapsed” across tissues within the same organ according to the GTEx label) for each gene.

Dimensionality reduction: We applied dimension reduction methods to the matrix of asymmetric coupling values (gene per row and gametolog pair–tissue combination per column). Dimension reduction was performed using Uniform Manifold Approximation and Projection (UMAP) via the umap function in the R package umap [82] with default parameters and the ‘manhattan’ metric. We also provide t-SNE and PCA plots in the supplement (using the Rtsne function (perplexity=30) in the R package Rtsne [83] and the prcomp function in the R package stats).

Variance partitioning: We performed row-wise variance partitioning on the matrix of asymmetric coupling values (gene per row and gametolog pair–tissue combination per column) using the fitExtractVarPart-Model and plotVarPart functions in the R package variancePartition [84]. This function allowed us to fit a linear mixed model for each gene to estimate contributions from multiple sources of variation while simultaneously correcting for all other variables. We modeled the asymmetric coupling values of N = 7,919 genes (with asymmetric coupling values to all pairs-tissues) as a function of separate terms for gametolog pair and tissue. Categorical terms (i.e., pair and tissue) were modeled as random effects, as recommended by the package’s creator [84]. We then extracted and visualized the fraction of variance explained by each term, in addition to the residual variance.

ANOVA: For each gene, we modeled variation in its asymmetric coupling values across gametolog pair–tissue combinations as a function of separate terms for gametolog pair and tissue using ANOVA (aov function in the R package stats). P-values were adjusted across genes using Benjamini-Hochenberg [77].

### Clustering and functional enrichment of genes with asymmetric coupling patterns

We estimated the Spearman correlations across groups of genes whose asymmetric coupling values were predicted (in ANOVA, see above) by gametolog pair only, tissue only, or both (using the cor.pairs function in the R package dcanr) [85] and then created distance matrices (using the as.dist function in the R package stats). For each gene set, we identified the optimal number of clusters using the NbClust function in the R package NbClust [86] with the following metrics: min.nc=2, max.nc=15, method=“average”, index = “dunn”. For the primary analysis of N=10,943 “pair only” genes, we retained clusters with >10 genes. This yielded 6 gene clusters (results shown in Figure 4H). The average asymmetric coupling values across genes in each cluster (for each tissue or gametolog pair) were visualized and the significance of this average value was obtained via permutation: 1) for each cluster we sampled N genes from the entire set (equal to number of genes in cluster) and estimated the mean asymmetric coupling values (per pair or tissue); 2) we repeated this procedure 100x (for computational tractability) to obtain a distribution of values; and 3) we compared the observed value to the distribution and defined significant observations as those where the absolute observed value is greater than 95% of the absolute permuted values (after correcting for multiple comparisons using Benjamini-Hochberg). Functional enrichments for each cluster were obtained using the enrichGO function in the R package clusterprofiler [87] (minGSSize = 5, maxGSSize = 500; ont = “CC” for cellular compartments or “BP” for biological processes). Core/hub genes were identified for each cluster by: 1) estimating Spearman’s rank order correlations for the asymmetric coupling values across all genes in a given cluster; ii) calculating an average correlation per gene; and iii) extracting the genes both with the highest average correlation values and associations with the biological function highlighted in the manuscript (e.g., translation, synapse, etc.).

### Estimating sex-biased gene expression

For analyses of sex-biased gene expression, we removed sex-specific or sex-differentiated tissues, resulting in N = 14,109 samples (4,543 females, 9,566 males) across N=40 tissues. The following statistical approach is similar to that of Oliva and colleagues [47]. In particular, we first estimated sex differences in expression for each tissue using the voom-limma approach [75]: the normalized expression level of each gene (see above) was modeled as a function of sex, age, RIN, ischemic time, and intronic rate using the lmFit function (R package limma), and sex effects were estimated using the makeContrasts, contrasts.fit, and eBayes functions (R package limma) [75]. We then applied multivariate adaptive shrinkage to the outputs from the voom-limma models described above (i.e., for each region, the per gene *β* and their standard errors) using the R package mashr [88]. For missing data, *β* were set to 0 and standard errors were set to 100 (as recommended by the mashr package’s creators). We first selected strong signals by running a condition-by-condition (1by1) analysis on all the data (mash_1by1 function) and extracting those results with local false sign rate (LFSR) < 0.01 in any condition. Specifically, this analysis runs ash in the R package ashr [89] on the data from each condition, an Empirical Bayes approach to FDR analysis that incorporates effect size estimates and standard errors, and assumes the distribution of the actual effects is unimodal, with a mode at 0. We also generated a random subset of the data (50% of expressed genes), computed a list of canonical covariance matrices (cov_canonical function), and used these data and matrices to estimate the correlation structure in the null tests (estimate_null_correlation function). We then set up the main data objects (i.e., “strong” and “random”) with this correlation structure in place (mash_set_data function). We used the strong tests to set up data-driven covariances by performing PCA on the data (using 5 PCs; cov_pca function) and using the resulting 5 candidate covariance matrices to initialize and perform “extreme deconvolution” (cov_ed function) [90]. We then estimated canonical covariances from the random tests and then fit mash to the random tests using both data-driven and canonical covariances. We extracted the fitted g mixture from this model and specified this mixture model when fitting mash to the strong tests.

We compared the overall (expression weighted) asymmetric coupling values (per gene and tissue; see above) to sex effects on gene expression (per gene and tissue; using *β* from the mashr analysis) by estimating Spearman’s rank order correlation (cor.test function in the R package stats) and running Fisher’s exact test (fisher.test function in the R package stats; alternative = ‘greater’) across N = 23,053 observations (N = 7,690 unique genes) with significant sex-bias (LFSR < 0.05) and overall asymmetric coupling (summed CLIP: p_adj_ < 0.05) in a given tissue (excluding Y chromosome genes) (results shown in Figure 5A).

### Overlap between asymmetric coupling and sex-biased co-expression

To test whether X- or Y-biased coupling was associated with female- or male-biased co-expression, respectively, we obtained lists of genes with female- or male-biased co-expression for N=22 tissues from Hartman and colleagues [51]. We limited our analysis to autosomal genes and tested whether X- or Y-biased gene sets (i.e., all genes with positive or negative overall asymmetric coupling values, respectively) were enriched for genes with female- or male-biased co-expression (i.e., 4 enrichment tests). We performed these enrichments both across and within tissues using Fisher’s exact tests (fisher.test in the R package stats; alternative = ‘greater’). P-values were adjusted across tissues and within comparisons using Benjamini-Hochenberg [77] (results shown in Figure 5B).

### Sex differences in co-expression between X-biased genes and Y-biased genes

To test whether males and females differ in the average magnitude of co-expression between X-biased genes and Y-biased genes, we defined X-biased or Y-biased gene sets (within each tissue) as those with significant overall asymmetric coupling (summed CLIP p_adj_<0.05, see above) and positive or negative t-values, respectively. After centering and scaling each tissue-specific male and female co-expression matrix (mean=0, sd=1) (scale function in the R package base), we subsetted the rows to Y-biased genes and the columns to X-biased genes obtained for the relevant tissue (and also to genes expressed in both males and females). We then averaged all co-expression values for males and females in each tissue, and tested for sex differences across tissues using a paired t-test (t.test function in the R package stats) (results shown in Figure 5C).

### Analysis of ASD risk genes implicated by rare variant analysis

We considered N = 102 ASD risk genes identified by Satterstrom and colleagues [58] that were partitioned into functional categories by the authors using gene ontology and literature searches (results shown in Figures 5D-F). Categories include gene expression regulation (N = 58), neuronal communication (N = 24), and genes in “other” categories (N = 20). We detected almost all risk genes (101/102) in at least one GTEx brain tissue (N = 13 tissues) – only the cerebellum-specific transcription factor *PAX5* was not detectably expressed. To formally test for associations between ASD gene sets and asymmetric coupling within each brain tissue, we: i) performed chi-squared tests on contingency tables of genes categorized by asymmetric coupling (X-biased, Y-biased, or unbiased) and ASD association/function (GER, NC, other); ii) tested whether genes in different ASD functional groups exhibited different mean asymmetric coupling values (regardless of CLIP p_adj_ value) using ANOVA and Tukey’s HSD tests; and iii) compared the observed mean asymmetric coupling values (regardless of CLIP p_adj_ value) and counts of asymmetrically coupled genes (p_adj_ < 0.05) within each ASD functional group to null distributions. Null comparisons were performed by: i) resampling N genes (equal to the number of genes within a given ASD gene set in a given tissue) – with each resampled gene matched to a gene in the original set according to their mean decile of expression in that tissue; ii) calculating the number of significantly X-biased and Y-biased genes; iii) estimating the mean asymmetric coupling value across genes; iv) repeating steps i-iii 1000x to create distributions of gene counts (from ii) and mean asymmetric coupling values (from iii); and v) comparing the observed values to those derived from the null distributions.

## Supplementary Text

### Asymmetric coupling to X vs. Y gametologs intersects with genetic risk for a sex-biased disorder

GER genes were X-biased in more brain areas (relative to genes in other functional categories; mean [range] # of regions across genes in each set: GER = 3.53 [1-10], other = 2.5 [1-4]) and NC genes showed Y-biased coupling in more brain areas (relative to genes in other functional categories; mean [range] # of regions across genes in each set: NC = 3.57 [1-7], GER = 2.15 [1-6], other = 1.89 [1-3]) (Table S23). GER genes tended to overlap with a higher proportion of X-biased genes and exhibit higher mean X-biased coupling values relative to: i) other functional categories (GER genes exhibit the highest proportion of X-biased genes in 10/10 brain tissues with >=1 asymmetrically coupled gene CLIP p_adj_ < 0.05; chi-squared tests: p_adj_ < 0.05 in 6/10; post-hoc Fisher’s exact tests [p_adj_ < 0.05]: GER > NC in 6/10, GER > non-ASD in 4/10, GER > other in 2/10) (average of mean asymmetric coupling values across tissues: GER = 0.03, non-ASD = −0.003, other = −0.02, NC = −0.05; ANOVA p_adj_ < 0.05 in 12/13 tissues; Tukey’s HSD [p_adj_ < 0.05]: GER > NC in 9/13, GER > non-ASD in 5/13, GER > other in 3/13) (Figure 5E; Tables S23, S24); and ii) null distributions (matched by mean expression decile) (GER genes overlap with a higher number of X-biased genes [pnull-adj < 0.05] in 4/10 tissues; GER genes exhibit higher mean X-biased coupling values [pnull-adj < 0.05] in 7/13) (Tables S23, S24). On the other hand, NC genes tend to overlap with a higher proportion of Y-biased genes and exhibit higher mean Y-biased coupling values relative to: i) other functional categories (NC genes exhibit the highest proportion of Y-biased genes in 6/10 tissues; chi-squared tests: p_adj_ < 0.05 in 6/10; post-hoc Fisher’s exact tests [p_adj_ < 0.05]: NC > GER in 6/10, NC > non-ASD in 4/10, NC > other in 2/10) (average of mean asymmetric coupling values across brain tissues: NC = −0.05, other = −0.02, non-ASD = −0.003, GER = 0.03; ANOVA p_adj_ < 0.05 in 12/13 tissues; Tukey’s HSD [p_adj_ < 0.05]: NC < GER in 9/13, NC < non-ASD in 8/13) (Figure 5E; Tables S23, S24); and ii) null distributions (matched by mean expression decile) (NC genes overlap with a higher number of Y-biased genes [pnull-adj < 0.05] in 4/10 tissues; NC genes exhibit higher mean Y-biased coupling values [pnull-adj < 0.05] in 9/13 tissues) (Tables S23, S24).

## Supplementary Figures

Figure S1 | Methods diagram. a. Bottom portion of Figure 1. b. Methods for estimating the significance of asymmetric coupling (overall and per pair) using the CLIP method. c. method for calculating overall asymmetric coupling.

Figure S2 | Hierarchical clustering dendrograms of tissues for between sex CFD (aCFDXXF–XYM) (a), within-pair absolute CFD (aCFDXM–YM) (b), and within-pair singed CFD (sCFDXM–YM) (c). Red values are AU (Approximately Unbiased) p-values, and green values are BP (Bootstrap Probability) values (see Methods).

Figure S3 | Between-sex signed co-expression fingerprint divergence (sCFDXXF–XYM) modeled as a function of within-pair signed co-expression fingerprint divergence in males (sCFDXM–YM). Regression lines are provided for each tissue (see legend) and across all data points (black line, confidence interval is shaded).

Figure S4 | Correlations across regulatory structural dissimilarity measures. Darker blue = more positive correlation; darker red = more negative correlation.

Figure S5 | a. X–Y co-expression values in the current study vs. those estimated by Godfrey and colleagues [20]. We recomputed X–Y co-expression values to be comparable to our estimates of functional divergence (aCFDXXF–XYM and sCFDXXF–XYM in Figures S5B and S5C). Differences between X–Y co-expression values across studies are the result of methodological differences, as our study: i) used a newer version of the GTEx data (v8 vs. v6); ii) removed age effects when calculating adjusted expression levels (Methods); and iii) normalized co-expression values by expression level using spatial quantile normalization (Methods). Regression lines are provided for each tissue (see legend) and across all data points (black line, confidence interval is shaded). These values are highly similar (*ρ* = 0.735, p < 2.2e-16; linear model: slope = 0.585; p < 2.2e-16) across pair–tissue combinations. b. aCFDXXF–XYM vs. X–Y member co-expression. Regression lines are provided for each tissue (see legend) and across all data points (black line, confidence interval is shaded). Absolute within-pair divergence (aCFDXM–YM) necessarily exhibits a strong negative correlation with the level of co-expression between the X and Y members themselves (within each pair) (*ρ* = −0.80, p < 2.2e-16). c. sCFDXXF–XYM vs. X–Y member co-expression. Regression lines are provided for each tissue (see legend) and across all data points (black line, confidence interval is shaded). Note that relative to aCFDXXF–XYM (Figure S5B), this relationship is relatively muted for signed divergence (sCFDXM–YM) (*ρ* = −0.24, p < 7.84e-08). d. aCFDXM–YM vs. the median Y/X expression ratio. Regression lines are provided for each tissue (see legend) and across all data points (black line, confidence interval is shaded). e. sCFDXM–YM vs. the median Y/X expression ratio. Regression lines are provided for each tissue (see legend) and across all data points (black line, confidence interval is shaded). f. Differences in X–Y expression variance vs. differences in X–Y mean expression. Regression lines are provided for each tissue (see legend) and across all data points (black line, confidence interval is shaded). g. Figure S5D excluding *TMSB4X/Y*. Regression lines are provided for each tissue (see legend) and across all data points (black line, confidence interval is shaded). h. Figure S5E excluding *TMSB4X/Y*. Regression lines are provided for each tissue (see legend) and across all data points (black line, confidence interval is shaded). i. Figure S5F excluding *TMSB4X/Y*. Regression lines are provided for each tissue (see legend) and across all data points (black line, confidence interval is shaded). j. Y/X expression ratios (log2) across all pair–tissue combinations. Purple = Y > X expression. Red = X > Y expression.

Figure S6 | a. Figure 4D excluding tissue-specific genes. Color indicates direction of coupling (see Figure 4B legend). Note that although the modal number of tissues in which genes showed significant X- or Y-biased coupling was one (Figure 4D), the current plot suggests that this pattern is driven by genes with tissue-specific expression. b. Count of genes with significant asymmetric coupling (CLIP p_adj_ < 0.05) to a given gametolog pair in a maximum of N tissues. Color indicates direction of coupling (see Figure 4B legend). c. Figure S6B excluding tissue-specific genes. Color indicates direction of coupling (see Figure 4B legend). d. Count of genes with significant asymmetric coupling (CLIP p_adj_ < 0.05) with a maximum of N pairs within a given tissue. Color indicates direction of coupling (see Figure 4B legend). Note that, while the modal number of gametolog pairs that genes were asymmetrically coupled to (across all tissues) was eight (Figure 4D), the modal number of pairs within a given tissue was only two (shown here). e. Number of genes with significant (p_adj_ < 0.05) asymmetric coupling for each pair–tissue using the subsamples (N = 66 males, see Methods) vs. the entire sample (*ρ* = 0.61, p < 2.2e-16). f. Number of genes with significant (p_adj_ < 0.05) asymmetric coupling for each pair using the subsamples (N = 66 males, see Methods) vs. the entire sample (*ρ* = 0.97, p = 2.767e-10). g. Number of genes with Methods (p_adj_ < 0.05) asymmetric coupling for each tissue using the subsamples (N = 66 males, see Methods) vs. the entire sample (*ρ* = 0.39, p = 0.01).

Figure S7 | a. Boxplots of asymmetric coupling values among autosomal, X chromosome, and Y chromosome genes with significant asymmetric coupling (CLIP p_adj_ < 0.05). Values are shown for each gametolog pair. b. Boxplots of asymmetric coupling values among autosomal, X chromosome, and Y chromosome genes with significant asymmetric coupling (CLIP p_adj_ < 0.05). Values are shown for each tissue. Non-gametolog Y chromosome genes show strong Y-biased coupling in the testes (e.g., *HSFY1/2, TSPY1/4/9*) and stomach (*DAZ1/4*).

Figure S8 | a. PCA of asymmetric coupling values across all pairs–tissues (represented by each point, see legend). b. tSNE of asymmetric coupling values across all pairs–tissues (represented by each point, see legend). c. UMAP of asymmetric coupling values across tall pairs–tissues (represented by each point, see legend). d. PCA of asymmetric coupling values across all pairs–tissues (represented by each point, see legend). Similar tissues have been collapsed (see Methods). e. tSNE of asymmetric coupling values across all pairs–tissues (represented by each point, see legend). Similar tissues have been collapsed (see Methods). f. UMAP of asymmetric coupling values across all pairs–tissues (represented by each point, see legend). Similar tissues have been collapsed (see Methods).

Figure S9 | a. First panel of Figure 4H. Mean asymmetric coupling values with each gametolog pair for genes in each cluster. N = 8 clusters were derived from N = 10,943 genes whose asymmetric coupling values were predicted by gametolog pair only (N = 2 clusters with < 10 genes were removed). Clusters and pairs are ordered according to hierarchical clustering. Asterisks indicate p_adj_<0.05 (vs. null, see Methods). b. First panel of Figure 4H, but with mean sCFDXM–YM values shown for each tissue. c. First panel of Figure 4H, but with pairs ranked within each cluster according to their mean sCFDXM–YM values. d. Similar to the first panel of Figure 4H, but for genes whose expression is predicted by tissue only (see Table S13, Column L). Mean asymmetric coupling values in each tissue for genes in each cluster. N = 5 clusters were derived from N = 66 genes whose asymmetric coupling values were predicted by tissue only. Clusters and pairs are ordered according to hierarchical clustering. Asterisks indicate p_adj_<0.05 (vs. null, see Methods). e. Figure S8D, but with mean sCFDXM–YM values shown for each pair. f. Figure S8D, but with tissues ranked within each cluster according to their mean sCFDXM–YM values. g. Similar to the second panel of Figure 4H, but for genes whose expression is predicted by tissue only (see Table S13, Column L). Top enriched biological processes and associated -log10(p-values) for each cluster. Top 4 categories are shown for each cluster. h. Similar to the third panel of Figure 4H, but for genes whose expression is predicted by tissue only (see Table S13, Column L). Top enriched cellular compartments and associated -log10(p-values) for each cluster. Top 4 categories are shown for each cluster. i. Similar to the first panel of Figure 4H, but for genes whose expression is predicted by both pair and tissue (see Table S13, Column M). Mean asymmetric coupling values for each pair (top) or tissue (bottom) for genes in each cluster. N = 8 clusters were derived from N = 1,333 genes whose asymmetric coupling values were predicted by pair and tissue. Clusters and pairs are ordered according to hierarchical clustering. Asterisks indicate p_adj_<0.05 (vs. null, see Methods). j. Figure S8I, but with mean sCFDXM–YM values shown for each tissue (top) and pair (bottom). k. Similar to the second panel of Figure 4H, but for genes whose expression is predicted by pair and tissue (see Table S13, Column M). Top enriched biological processes and associated -log10(p-values) for each cluster. Top 4 categories are shown for each cluster. l. Similar to the third panel of Figure 4H, but for genes whose expression is predicted by pair and tissue (see Table S13, Column M). Top enriched cellular compartments and associated -log10(p-values) for each cluster. Top 4 categories are shown for each cluster. Figure S10 | Figure 5A with tissue specific regression lines.

## Supplementary Tables

Table S1 | Tissue metadata, including the number of male and female samples and number of genes expressed per tissue.

Table S2 | Between-sex and within-pair CFD values (and associated confidence intervals and p-values) for each gametolog pair and tissue.

Table S3 | Mean between-sex CFD values (and associated p-values) per tissue.

Table S4 | Correlations between between-sex and within-pair CFD values for each tissue (for both absolute and signed CFD values). Significant p_adj_ < 0.05 are in bold.

Table S5 | Dissimilarity measures for each gametolog pair, associated evolutionary strata, and correlations (*ρ* and p-values) between regulatory/structural dissimilarity measures and mean within-pair absolute functional divergence (mean aCFDXM–YM).

Table S6 | Correlations across promoter, coding, and protein dissimilarity measures (*ρ* and p-values).

Table S7 | Asymmetric coupling values for each expressed gene (listed in Column 1) for each gametolog pair–tissue combination (listed in column headings).

Table S8 | Asymmetric coupling CLIP adjusted p-values for each expressed gene (listed in Column 1) for each pair–tissue combination (listed in column headings).

Table S9 | Asymmetric coupling values for each expressed gene (listed in Column 1) for each pair–tissue combination (listed in column headings) (subsampled to N=66 males per tissue).

Table S10 | Asymmetric coupling CLIP adjusted p-values for each expressed gene (listed in Column 1) for each pair–tissue combination (listed in column headings) (subsampled to N=66 males per tissue).

Table S11 | Expression-weighted average asymmetric coupling values and associated CLIP p-values for each expressed gene (listed in Column 1) in each tissue (listed in Column 2).

Table S12 | Results of variance partitioning analysis (for N = 7,919 genes). Values represent the proportion of variance in expression of each gene (listed in Column A) that is explained by pair (Columns C or G), tissue (Columns D or H), or residual variation (Columns E or I). Results are shown for analyses using all tissues (Columns C-E) and using collapsed tissues (Columns G-I).

Table S13 | Results of ANOVA models (asymmetric coupling gametolog pair + tissue) including F-values, p-values, and adjusted p-values. Results are shown for analyses using all tissues (Columns C-M) and using collapsed tissues (Columns O-U). Cluster assignments are shown for genes whose asymmetric coupling is predicted by gametolog pair only (Column K), tissue only (Column L), or both (Column M).

Table S14 | GO (biological process) enrichment results for each cluster (listed in Column 1) derived from genes whose expression is predicted by gametolog pair only (see Table S13, Column K).

Table S15 | GO (cellular compartment) enrichment results for each cluster (listed in Column 1) derived from genes whose expression is predicted by gametolog pair (see Table S13, Column K).

Table S16 | GO (biological process) enrichment results for each cluster (listed in Column 1) derived from genes whose expression is predicted by tissue only (see Table S13, Column L).

Table S17 | GO (cellular compartment) enrichment results for each cluster (listed in Column 1) derived from genes whose expression is predicted by tissue only (see Table S13, Column L).

Table S18 | GO (biological process) enrichment results for each cluster (listed in Column 1) derived from genes whose expression is predicted by both gametolog pair and tissue (see Table S13, Column M).

Table S19 | GO (cellular compartment) enrichment results for each cluster (listed in Column 1) derived from genes whose expression is predicted by both gametolog pair and tissue (see Table S13, Column M).

Table S20 | Sex-biased gene expression results (from mashr; *β* and LFSRs) for each expressed gene (Column 1) in each tissue (N=40; Column 2).

Table S21 | Tissue specific correlations (number of genes, *ρ*, and p-values) between asymmetric coupling and sex-biased gene expression. Significant p_adj_ < 0.05 are in bold.

Table S22 | Tissue-specific enrichment results (Fisher’s exact tests) between genes with X- or Y-biased coupling and genes with sex-biased co-expression. OR = Odds ratio. Significant p_adj_ < 0.05 are in bold.

Table S23 | N = 101 ASD risk genes (from Satterstrom and colleagues) detected in the GTEx data. Tissue-specific asymmetric coupling values (and CLIP p-values) are shown for N = 13 brain tissues.

Table S24 | Enrichment tests linking ASD risk gene sets to asymmetric coupling.

## Acknowledgements

The Genotype-Tissue Expression (GTEx) Project was supported by the Common Fund of the Office of the Director of the National Institutes of Health, and by NCI, NHGRI, NHLBI, NIDA, NIMH, and NINDS. The data used for the analyses described in this manuscript were obtained from dbGaP accession number phs000424/GRU. This research was supported (in part) by the Intramural Research Program of the NIMH (1ZIAMH002949-03).

---

Note 1: Throughout, we use the term “sex” to refer to a label that people are assigned at birth based on their anatomy, which often corresponds with one of two sex chromosome complements. We use the phrase “sex differences” to refer to group-level average differences between individuals with “typical” (of the majority) anatomical and sex chromosome complements (i.e., XY males with testes, XX females with ovaries), although we acknowledge that these criteria are not confirmed in many of the human studies discussed here (which instead rely on self identification). In our analyses of the GTEx data, we defined female and male donors exclusively on the basis of their sex chromosomes, XX or XY, respectively. Across all individuals, karyotypic and self identified sex were in agreement [41]. We acknowledge that while most people are categorized as female or male at birth, sex is not strictly binary. Intersex people represent 1% of the population and exhibit an abundance of variation across sex chromosome combinations, sex hormone concentrations and receptors, and bodily phenotypes. We also distinguish sex from gender, a culturally defined and malleable concept. A person’s gender need not align with their assigned sex, and since an individual’s experiences in society can be impacted by their perceived gender, these biological and psychosocial influences are difficult (if not impossible) to disentangle in humans.

